# Systemic migrations in enucleated cells

**DOI:** 10.1101/2024.05.19.594852

**Authors:** Ildefonso M. De la Fuente, Jose Carrasco-Pujante, Borja Camino-Pontes, Maria Fedetz, Carlos Bringas, Alberto Pérez-Samartín, Gorka Pérez-Yarza, José I. López, Jesus M Cortes, Iker Malaina

**Author notes:** **Corresponding Author:** Correspondence and requests for materials should be addressed to Dr. Ildefonso M. de la Fuente. Instituto CEBAS-CSIC. Campus Universitario de Espinardo. Espinardo. 30100 Murcia. ESPAÑA. /. Equal contribution (Jesus M Cortes, Iker Malaina).

## Abstract

Directional locomotion is a fundamental characteristic of many cells with great relevance in essential physiological and human pathological processes. For decades, research efforts have focused on studying specific individual processes and their corresponding biomolecular components involved in cellular locomotion movements. However, the notion that migratory displacements are functionally integrated and regulated at the systemic level has never been recognized. Recently, we have shown that locomotion movements correspond to an emergent systemic behavior which depends on a complex integrated self-organized system carefully regulated at a global cellular level. Here, to study the forces driving the locomotion movement of the cell, and to corroborate the thesis on the systemic character of the cellular migratory responses we have carried out an extensive study of locomotion movements with enucleated cells (cytoplasts) belonging to *Amoeba proteus*. The migratory behavior of both enucleated and non-enucleated cells has been individually studied in four different scenarios: in absence of stimuli, under a galvanotactic field, in chemotactic gradient, and under very complex conditions such as simultaneous galvanotactic and chemotactic stimuli. All these experimental displacement trajectories, obtained on flat two-dimensional surfaces, have been analyzed using advanced non-linear quantitative approaches. Our results show that both non-enucleated amoebas and cytoplasts display the same complex kind of dynamic structure in their migratory trajectories. The locomotion displacements of enucleated cells are regulated by complex self-organized integrative dynamics, modulated at a global-systemic level which seems to depend on the cooperative interaction of most, if not all, molecular components of cells.

## Introduction

For a wide range of cells, from prokaryotes to eukaryotes, locomotion movements correspond to a fundamental complex behavior of unicellular organisms endowed with migratory properties.

Free-living cells efficiently explore their external environment to find food, adjusting with precision their trajectory and speed while avoiding predators and adverse conditions. This ability to directionally move is critical for life in Metazoan organisms. Thus, migratory displacements play a central role in essential biological processes, such as embryogenesis (creation of the endodermal, mesodermal and ectodermal layers during gastrulation), organogenesis, neural development, wound healing (e.g., somatic stem cell types for tissue repair, as well as mesenchymal and hematopoietic stem cells, display remarkable migratory capabilities, mobilizing to injury sites to participate in their regeneration), angiogenesis, and immune responses (SenGupta et al., 2021).

Furthermore, misguided cellular locomotion responses in humans play a significant role in numerous pathologies. These include atherosclerosis (Charo and Taubman, 2004), the progression of metastatic tumors (Eccles, 2005; Stuelten et al., 2018; De la Fuente and López, 2020; Novikov et al., 2021), progeria (Chang et al., 2019), osteoarthritis (Qu et al., 2019), fibrosis (Glass et al., 2020; Tisler et al., 2020; Hewitt et al., 2023), auditory impairments (Mohamed et al., 2023), chronic obstructive pulmonary disease (Wu et al., 2020), immune-related actinopathies (Kamnev et al., 2021), endometriosis (Jiang et al., 2015; Choi et al., 2019), multiple sclerosis (Al Abadey et al., 2022; Rodriguez-Mogeda et al., 2022), rheumatoid arthritis (Zhou et al., 2021; Kotschenreuther et al., 2022), diabetic wound (Pawar et al., 2021), psoriasis (Cumberbatch et al., 2006; Eaton et al., 2018), asthma (Chang et al., 2011; Zhang et al., 2016; Salter et al., 2017), Crohn’s disease (Petagna et al., 2020; Mao et al., 2022), congenital brain disorders (Spalice et al., 2009; Klingler et al., 2021; Edey et al., 2023), and other types of immunodeficiencies (Janssen and Geha, 2019).

The important physiological processes in which cell migration is involved, as well as its related pathologies, make this complex cellular behavior a key issue in contemporary biology. Specifically, one of the central questions regarding cellular directed movement is whether this cellular property corresponds to a coordinated and integrated emerging process at the global cellular level.

Recently, an exhaustive study has shown that the locomotion movements seem to depend on a complex integrated self-organized system carefully regulated at global level, arising from the cooperative non-linear interaction of most, if not all, cellular components. Such emergent systemic property seems to be not found in partial mechanisms, in any specific molecular part, or individual physiological processes of the cell (De la Fuente et al., 2024).

Supporting this thesis, a notable number of experimental researches has shown that practically the majority of cellular physiological processes are involved in cell migration. In fact, in addition to the main components of cytoskeleton (actin microfilaments, microtubules, and intermediate filaments) energy is a vital factor in cell motility; the cytoskeleton converts chemical energy into mechanical forces requiring significant bioenergetic demands; hence, mitochondrial activity and the adenylate energy system (De la Fuente et al., 2014) are key regulators of directional motion (Fujisawa, 2023). The endoplasmic reticulum, a signaling organelle controlling multiple processes like calcium ions, also contributes to cell locomotion regulation (Sáez et al., 2014; Limia et al., 2019). Furthermore, cell polarity, influenced by centrosome positioning (Ueda et al., 1997; Zhang and Wang, 2017) and centrosomal orientation (Schmoranzer et al., 2009; Chang et al., 2015), is essential for directional motion. The Golgi apparatus, a key processing centre for proteins modified in the endoplasmic reticulum, modulates intracellular traffic processes, influencing cell motility (Mellor, 2004). The global nucleus structure, acts as a central mechanosensory system, being crucial for appropriate mechanical responses during cell migration (Fruleux and Hawkins, 2016; Long and Lammerding, 2021). Structural chromatin organization also plays a significant role in cellular migration (Gerlitz, 2020). In addition, numerous biochemical molecules and other physiological processes are involved in the directional movement of cells, including PI3K (Inoue and Meyer, 2008), focal adhesion proteins (Artemenko et al., 2014), actin-binding proteins (Lee and Dominguez, 2010), PLEKHG3 (Nguyen et al., 2016), p21-activated kinases (Kumar et al., 2009), SCAR/WAVE proteins (Pollitt and Insall, 2009), Arp2/3 complexes (Swaney and Li, 2016), mitogen-activated protein kinases (Huang et al., 2004), actin-capping protein (Ogienko et al., 2013), WASP family proteins (Veltman and Insall, 2010), phosphoinositide lipid molecules (Ling et al., 2006; Kakar et al., 2023; Thapa et al., 2023), TORC2/PKB pathway (Cai et al., 2010), and the Nck family of adaptor proteins (Rivera et al., 2006), among others.

These experimental evidences indicate the existence of complex functional and integrative physiological-biomolecular processes involved in migratory behaviors at a cellular global scale.

Here, to understand the forces driving the locomotion movement of the cell and to corroborate the thesis that the directional motility of cells is regulated by integrative processes operating at the systemic level, we have quantitatively analyzed the experimental movement trajectories of individual enucleated cells (enucleated by micromanipulation) belonging to *Amoeba proteus* using advanced non-linear physical-mathematical tools and computational methods. For such purpose, we have studied the essential characteristics of the locomotion movement in 200 non-enucleated amoebas and 200 cytoplasts (enucleated cells) on flat two-dimensional surfaces and under starvation conditions. The migratory behavior of both the enucleated and non-enucleated cells has been individually studied in four different scenarios: in absence of stimuli, under a galvanotactic field (the electric membrane potential of cells enables predators like amoebas to detect prey), in a chemotactic gradient (we have used a nFMLP peptide, which indicates to the amoebae the possible presence of food in their immediate environment), and under very complex conditions such as simultaneous galvanotactic and chemotactic stimuli.

Historically, it is known that individual cells enucleated by micromanipulation can survive for extended periods, with a lifespan of up to two weeks (Ord, 1968). Research confirmed the full viability of cells without nuclei by reintroducing the nucleus after a 12-day interval; notably, certain amoebas could restore their full physiological activity, including cell division and the establishment of stable cultures (Ord, 1968). It is noteworthy that this 12-day enucleation period significantly exceeds the typical 24-hour cell cycle of *Amoeba proteus*, which can vary with environmental conditions but generally persists in a controlled setting (Prescott, 1955).

Our results show that both non-enucleated amoebas and cytoplasts display the same kind of complex dynamic structure of migratory movements which exhibits significant non-trivial long-range correlations, pronounced anomalous super-diffusion, persistent effects with positive trend-reinforcing behaviour, structured move-step sequences with minimal entropy and high information content, and effective exploration of the external environment. This migration structure singularizes the way in which locomotion movements occur, and such characteristics correspond to critically self-organized systems.

The migratory trajectories of both non-enucleated and enucleated amoebas change continually since they are characterized by random magnitudes that vary over time, but nevertheless the cellular stochastic displacements shape a type of dynamic locomotion structure whose defining characteristics are unambiguously preserved in both enucleated and non-enucleated cells in all the conditions analyzed.

These results with enucleated cells corroborate our previous published results (De la Fuente et al., 2024) which showed that migratory responses seem to be regulated by functional integrative processes operating at systemic level.

In short, the locomotion trajectories of enucleated cells correspond to complex integrative dynamics, carefully regulated at a global-systemic level which seems to depend on the cooperative non-linear interaction of most, if not all, of the molecular components of cells.

## Results

To perform all our experiments, we used a special set-up consisting of two standard electrophoresis blocks (17.5 cm long), two agar bridges, a power supply, and a glass structure placed in the middle of the second electrophoresis block, where the amoebae were located (see *Materials and Methods* and *Supplementary Information*, Fig. S1). We plugged one electrophoresis block into a power supply unit and connected the other one to it with two agar bridges. This way, the anode and cathode did not touch the medium (Chalkley’s simplified medium (Korohoda et al., 2000)) where the amoebae swam. The glass structure made it possible to establish a laminar flux that let the electric current go through while also generating an nFMLP peptide gradient (see *Materials and Methods* and *Supplementary Information*, Figs. S1 and S2).

Before conducting each experiment, the amoebae underwent a 24-hour starvation period and, when necessary, were enucleated as described in *Materials and Methods* and Fig. 1*C* and *D*. We used a stereo microscope equipped with a digital camera to capture their individual migration patterns for 34’10’’. To reduce the influence of social interactions among the amoebae, each replication consisted of small groups of no more than 8 cells. The following basic experimental information data (BEID) parameters are provided for each scenario: “Nr” for the number of cells per replication, “Er” for the number of experimental replications, and “N” for the total number of cells. The recorded trajectories were then subjected to time series analysis using sophisticated non-linear dynamic techniques.

**Fig. 1.**
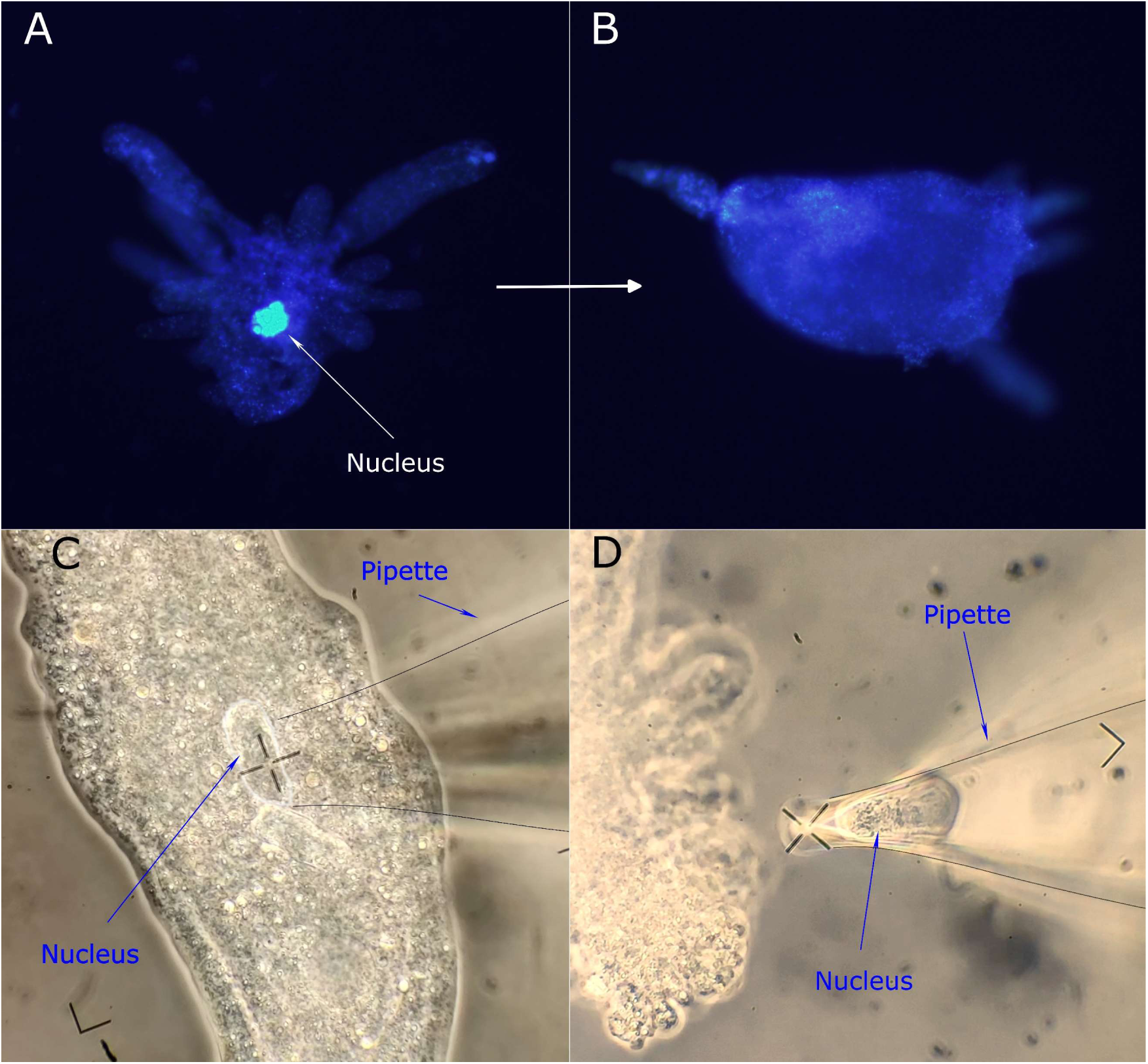
Cell enucleation procedure. *(A)* Fluorescent microscopy image of *Amoeba proteus* cell with its nucleus intact. *(B)* Fluorescent microscopy image of an enucleated cell (cytoplast). *(A and B)* Cytoplasts were stained with DAPI (1 μg/ml) after cell enucleation to confirm the absence of nucleus. *(C)* Enucleation process 1: extracting the nucleus with a glass enucleation pipette. *(D)* Enucleation process 2: enucleated cell (cytoplast), the extracted nucleus can be seen inside the enucleation pipette. See *Materials and Methods* for more details. Enucleation pipettes in *C* and *D* have been outlined with a thin black stroke to enhance visualization.

We recorded and analyzed the migratory movements of 400 amoebae (200 cells and 200 cytoplasts) belonging to the species *Amoeba proteus* under four experimental scenarios: without any stimulus, within an electric field, within a chemical gradient, and under simultaneous chemical and electrical stimuli.

### 1. Cell motility in the absence of external cues

The locomotion trajectories of 50 cells and 50 cytoplasts in a stimulus-free medium were monitored for 34 minutes and 10 seconds (Fig. 2*A* and*E*). Movement directionality was quantified by computing the displacement cosine for each trajectory (Fig. 2*A’* and *E’*). Values close to −1 imply a leftward bias, while values close to 1 indicate a rightward bias. Our analysis revealed that values spanned between −1 and 1, with a median/IQR of −0.05/1.35. Median/IQR values for each cell type were −0.28/1.28 (cells) and 0.09/1.40 (cytoplasts). These results suggest that amoebae moved randomly with no preferred direction in the absence of stimuli. BEID: N=50, Er=17, Nr=1-4 (cells); N=50, Er=34, Nr=1-4 (cytoplasts).

**Fig. 2.**
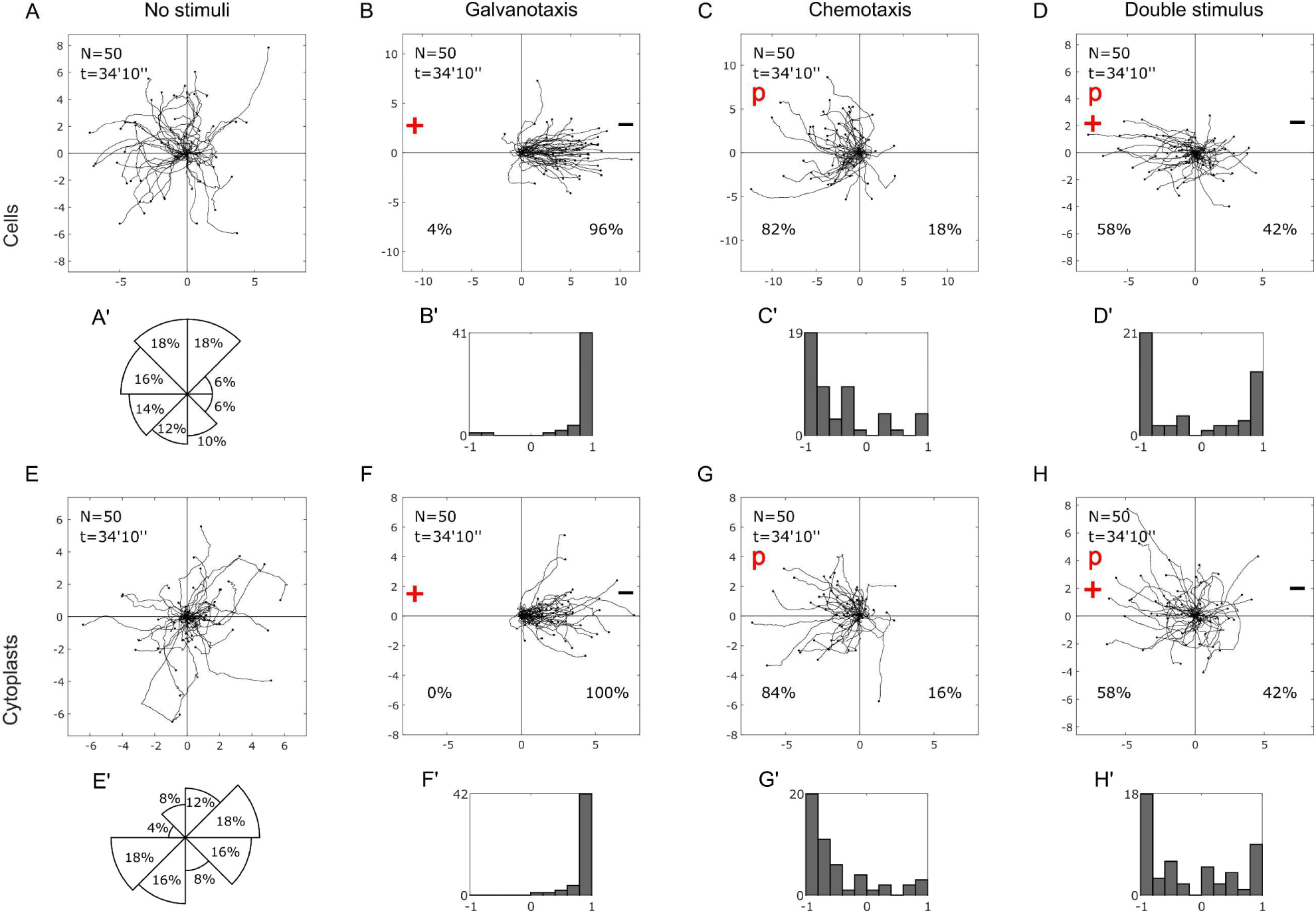
Cell and cytoplast motion under four experimental scenarios. *(A–H)* Migratory trajectories of *Amoeba proteus* cells (upper row) and cytoplasts (lower row) under four different experimental scenarios (*No stimuli*, *Galvanotaxis*, *Chemotaxis* and *Double stimulus*). Direction of cell *(A’)* and cytoplast *(E’)* displacements, displayed as the approximated percentage of trajectories that finished within each 45° sector of the experimental chamber, represented here as a polar chart. *(B’–D’* and *F’–H’)* displacement cosines corresponding to panels *(B–D)* and *(F–H)*. “N” total cell count; “t” experiment duration; “p” chemotactic peptide (nFMLP) location; “+” anode location; “−” cathode location. The starting point for each trajectory is located at the center of the plot, and the distance in mm is shown on both the x and y-axis.

### 2. Effects of an electric field on cell displacement

We recorded the trajectories of 50 cells and 50 cytoplasts migrating in the presence of a 300-600 mV/mm electric field (Fig. 2*B* and *F*). Amoebae overwhelmingly migrated towards the cathode, located on the right side of the set-up. The displacement angle cosines for all trajectories were calculated to assess movement directionality. Median/IQR values were 0.98/0.13 for cells, 0.96/0.12 for cytoplasts and 0.97/0.13 for all trajectories (Fig. 2*B’* and *F’*), confirming the emergence of a new behavior in the presence of an electric field. The Wilcoxon rank-sum test was used to compare values for the displacement cosines in the absence of stimuli and under an electric field. The results show that a new migration pattern with an extremely low chance to occur at random emerged in both cells and cytoplasts. (p-values: 10^−12^ and 10^−9^; Z: −7.08 and −5.98 for cells and cytoplasts, respectively). BEID: N=50, Er=11, Nr=4-8 (cells); N=50, Er=29, Nr=1-4 (cytoplasts).

### 3. Effects of a peptide chemical gradient on cell locomotion

We observed the migration trajectories of 50 cells and 50 cytoplasts under a chemotactic gradient of nFMLP peptide, which was applied on the left side of the experimental setup (Fig. 1*C* and *G*). Most of the amoebae (83%) moved towards the source of the peptide. We calculated the displacement angle cosines for each of the 100 trajectories, which ranged from −1 to 1. The median/IQR values were −0.66/0.65 (cells) and −0.71/0.57 (cytoplasts) and −0.70/0.65 for all trajectories, suggesting a consistent pattern of movement towards the peptide (Fig. 1*C’* and *G’*). We used the Wilcoxon rank-sum test to compare the cosine values in the presence of an electric field or a chemotactic gradient (p-values: 10^−13^ and 10^−15^; Z: 7.43 and 7.88 for cells and cytoplasts, respectively). These results indicated that the chemotactic gradient induced a distinct locomotion behavior from the one elicited by the presence of an electric field both in cells and cytoplasts. BEID: N=50, Er=12, Nr=3-7 (cells); N=50, Er=21, Nr=1-4 (cytoplasts).

### 4. Cell migration under combined electric and chemical stimuli

We analyzed the migratory trajectories of 100 amoebae (50 cells and 50 cytoplasts) under the simultaneous application of an electric field and a chemotactic gradient of nFMLP peptide. The peptide was located on the left side of the set-up (anode), while the right side was the cathode (Fig. 1*D* and *H*). In this condition, we observed that 42% of the amoebae moved towards the cathode, while 58% moved towards the peptide (anode). The median/IQR values of the displacement angle cosines for the 100 trajectories were −0.44/1.60. For cells and cytoplasts, the values were −0.4/1.76 and −0.55/1.41, respectively. These values indicated that the amoebae exhibited two distinct migratory behaviors, one towards the anode and one towards the cathode (Fig. 1*D’* and *H’*). The Wilcoxon rank-sum test revealed that these behaviors were significantly different for both cells (p-value=10^−9^; Z=5.98) and cytoplasts (p-value=10^−9^; Z=5.98). BEID: N=50, Er=13, Nr=3-4 (cells); N=50, Er=15, Nr=2-4 (cytoplasts).

### 5. Influence of long-range correlations on cellular locomotion

Complex systems exhibit systemic behavior that is characterized by the existence of long-range correlated dynamics (Prigogine, 1978). A widely used technique to detect these correlations in time series data (such as migratory patterns in this case) is the “root mean square fluctuation” (“rmsf”) analysis, a traditional approach in Statistical Mechanics that draws on the concepts proposed by Gibbs (Gibbs, 1902) and Einstein (Einstein, 1909).

A power-law relation of the form *F*(*l*) ∼ *l*^*a*^ can reveal long-range interdependence, where *l* is the number of steps. The fluctuation exponent *α* is around 0.5 for uncorrelated data, while *α* > 0.5 or *α* < 0.5 signify the existence of positive or negative long-range correlations respectively (see *Materials and Methods*). Fig. 3*A* shows an example of “rmsf” analysis for the locomotion patterns of a typical cell (taken from Scenario One) and cytoplast (taken from Scenario Three).

**Fig. 3.**
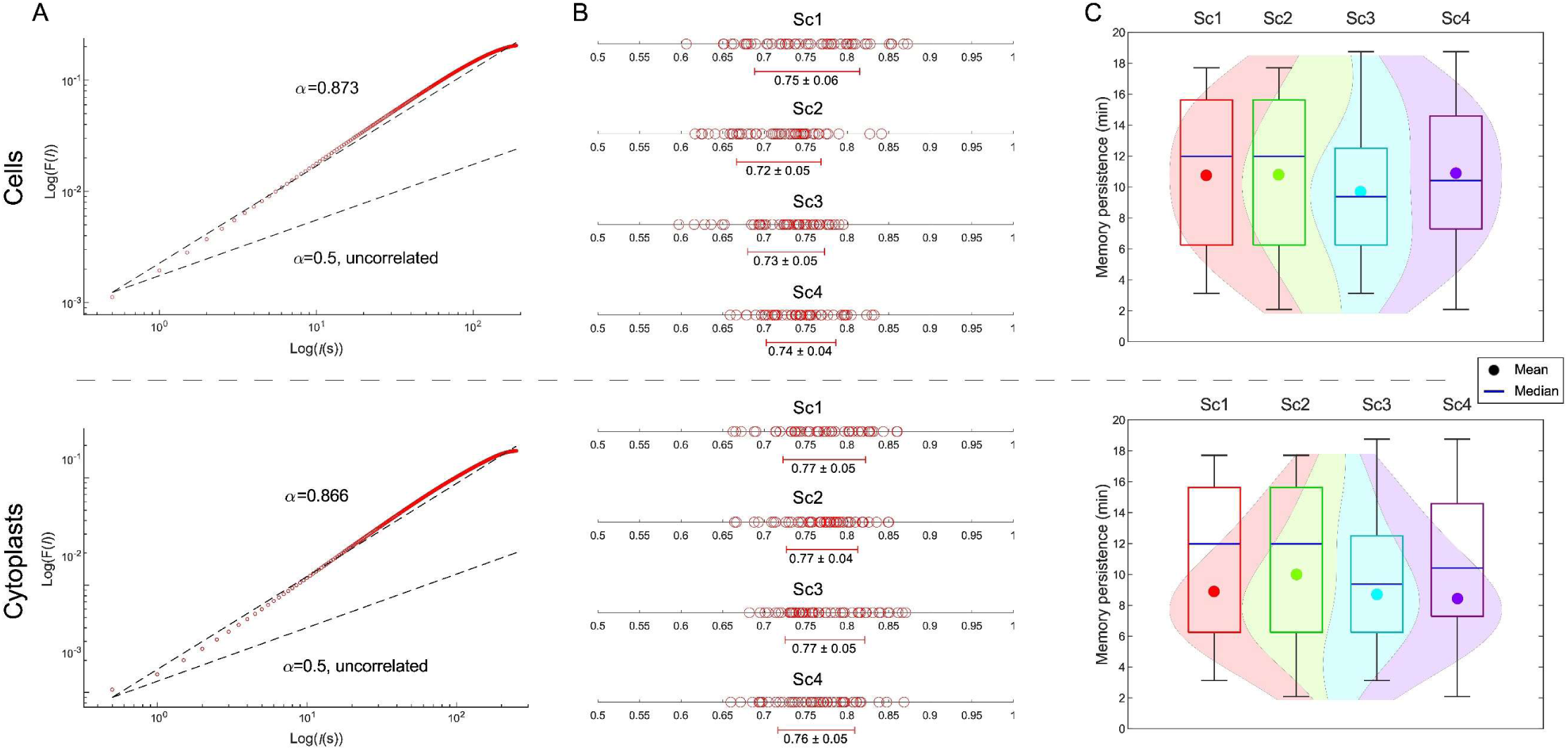
Move-steps of cellular migratory displacements show mutual interdependence over long periods of time. *(A)* Log-log plots of rmsf *F* versus *l* step for both a prototypical *Amoeba proteus* cell (upper panel) belonging to Scenario One (Sc1) and cytoplast (lower panel) belonging to Scenario Three (Sc3). *(B)* Linear graphs display the values (and global average ± SD) of experimental rmsf correlation coefficient *α* for all cells (upper panel) and cytoplasts (lower panel) under each experimental scenario. *(C)* Violin plots depict the distribution, mean, and median memory duration values for cells (upper panel) and cytoplasts (lower panel) in the four experimental scenarios. “Sc1” Scenario One (no stimuli); “Sc2” Scenario Two (galvanotaxis under an electric field); “Sc3” Scenario Three (chemotaxis along a peptide gradient) and “Sc4” Scenario Four (simultaneous galvanotactic and chemotactic cues).

To test the reliability of this technique, we shuffled all the 400 experimental trajectories (2 × 10^5^ shuffles per trajectory) and compared their scaling exponent *α* values to the original ones. Fig. 3*B* displays the “rmsf” analysis results of the 400 experimental amoeba trajectories (see *Supplementary Information*, Table S1 for more details). The cellular move-step migratory fluctuations of all the experimental trajectories showed non-trivial correlations. We observed that the “rmsf” scaling exponent *α* had a median/IQR of 0.74/0.07 for cells and 0.77/0.07 for cytoplasts. The “rmsf” analysis values of the total experimental trajectories ranged from 0.60 to 0.87, with a median/IQR of 0.75/0.07, while the *α* values of all shuffled trajectories ranged from 0.37 to 0.62, with a median/IQR of 0.47/0.07 (see *Supplementary Information*, Table S2 for more details). A Wilcoxon test comparing the experimental exponents to the shuffled ones revealed highly significant differences between our experimental and shuffled data (p-value≅0, Z=24.48), suggesting the unlikelihood of random occurrence of the observed long-range correlations in the experimental data.

We also measured the time span of the correlations regime and observed that all amoebae showed long-range coordination over periods varying from 2.08 to 18.75 minutes with a median/IQR of 9.38/7.29 minutes. These results imply strong dependences of previous movements lasting about 1125/875 (median/IQR) move-steps (Fig. 3*C* and *Supplementary Information*, Table S3). Cells and cytoplasts displayed long-range correlations with a median/IQR value of 10.42/8.33 and 8.33/5.21 minutes, respectively, which emphasizes the effect of prior trajectory values on each move-step. These findings indicate the existence of non-trivial long-range correlations in all displacement trajectories.

### 6. Efficient system dynamics in cellular directed movement

The strong anomalous dynamic of cell migration is another feature of cell locomotion. This is linked to anomalous super-diffusion, a complex process which is non-linear with respect to time and leads to effective directional trajectories at the system level (Faustino et al., 2007; Viswanathan et al., 2008). We calculated the Mean Square Displacement (MSD), a method developed by Einstein (Einstein, 1905) and von Smoluchowski (von Smoluchowski, 1906), to measure this dynamic property. This tool from Statistical Mechanics enables the quantification of the space covered by the amoebae during their movement. The anomalous diffusion exponent *β* is used in this method to indicate if normal (Brownian, *β* = 1) or anomalous diffusion (*β* ≠ 1) occurs (see *Materials and Methods*). Sub-diffusion and super-diffusion dynamics are associated with 0 < *β* < 1 and *β* > 1, respectively.

We performed an MSD analysis of the locomotion movements of a representative cell and cytoplast without stimuli, as shown in Fig. 4A. The MSD analysis of the 400 experimental cells (Fig. 4B, Fig. 4C and *Supplementary Information*, Table S4) revealed that almost all trajectories display strong anomalous migratory dynamics. The variable *β*, which describes the diffusion process, had a median/IQR value of 1.93/0.11 for cells and 1.91/0.11 for cytoplasts in the experimental trajectories. These values indicate an anomalous super-diffusive process, a complex behavior that seems to dominate the cell trajectories in both groups.

**Fig. 4.**
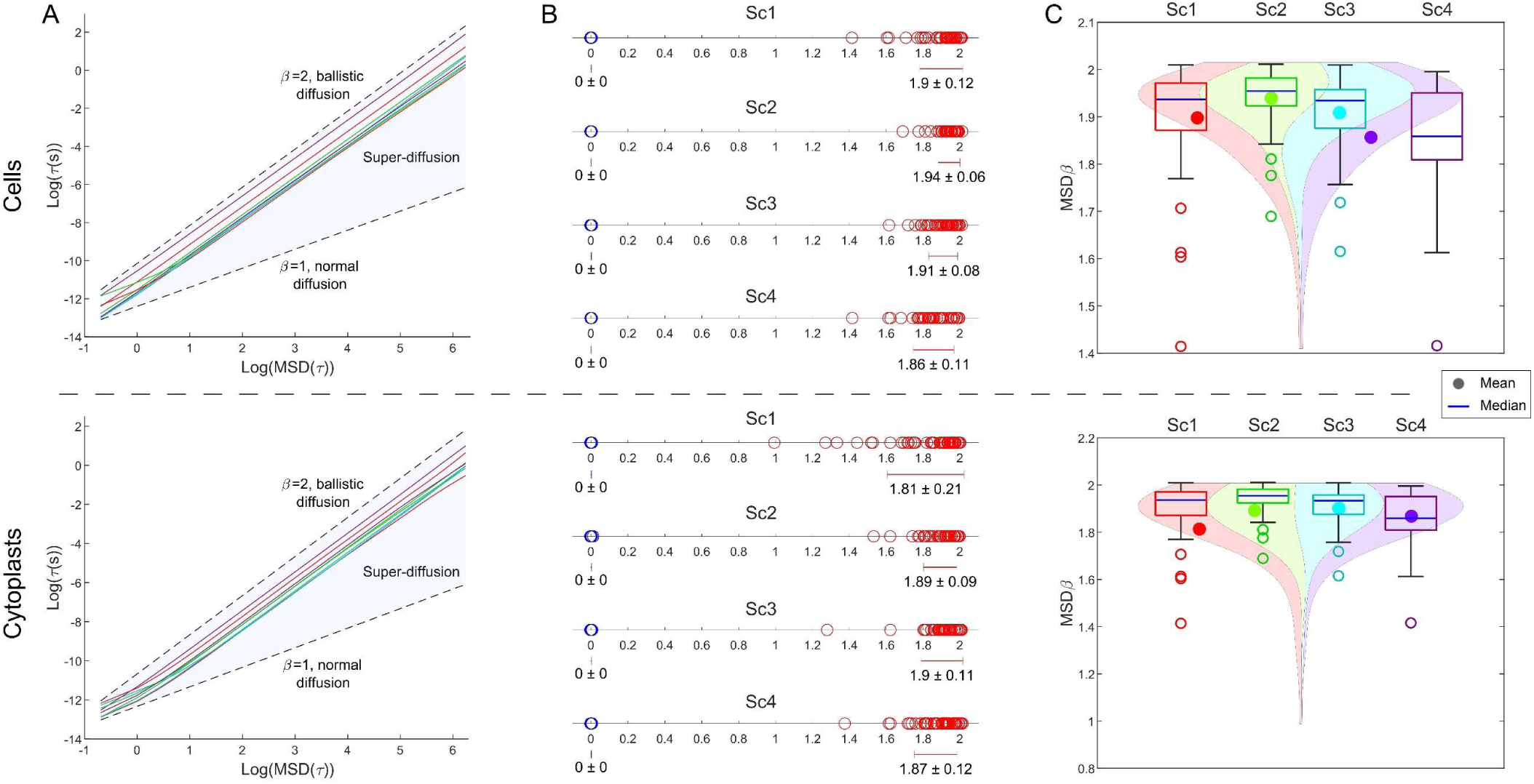
Strong anomalous migration patterns in cellular movement. *(A) β* values of the MSD for eight representative cells (upper panel) and cytoplasts (lower panel) belonging to Scenario One (Sc1). *β* = 1 indicates normal diffusion, *β* = 2 ballistic diffusion, and the shaded region that lies in between is indicative of super-diffusion, a multifaceted process exhibiting a pronounced non-linear correlation with time, encompassing all the observed values of the *β* exponent from the experimental trajectories. *(B)* Linear graphs illustrate all *β* exponent values (and their global average ± SD) for cells (upper panel) and cytoplasts (lower panel) under the four experimental scenarios, with experimental values in red and shuffled values in blue. By shuffling, the long-term correlation structure was eliminated, which resulted in a clear separation of the experimental and shuffled value distributions (p-value≅0, Z=24.48) across all cell types and experimental scenarios. *(C)* Violin plots show the estimated distribution, median and mean of the MSD *β* exponent values from all experimental cell (upper panel) and cytoplast (lower panel) trajectories. “Sc1” Scenario One (no stimuli); “Sc2” Scenario Two (galvanotaxis under an electric field); “Sc3” Scenario Three (chemotaxis along a peptide gradient) and “Sc4” Scenario Four (simultaneous galvanotactic and chemotactic cues).

Values of the anomalous diffusion exponent *β* for all experimental trajectories ranged from 0.99 to 2.01, with a median/IQR value of 1.92/0.12, while the shuffled trajectories had values from −0.00 to 0.01, with a median/IQR value of 10^−4^/0.00 (Fig. 4B and *Supplementary Information*, Table S5). A Wilcoxon test comparing experimental to shuffled anomalous diffusion exponents demonstrated that our results are highly improbable to occur by chance (p-value≅0, Z=24.48).

### 7. Unpredictability and information content in migratory trajectories

We used the Approximate Entropy (ApEn), a reliable estimate of the Kolmogorov–Sinai (K-S) entropy (Pincus, 1991; Delgado-Bonal and Marshak, 2019), to measure the information content within locomotion trajectories and shed light on the complex migratory behavior that results from the cellular system. Approximate Entropy (ApEn), introduced by Pincus et al. in 1991 (Pincus et al., 1991), serves as a statistical tool to quantify the regularity and complexity of fluctuations in time-series data. A series with intricate and challenging-to-anticipate behavior demonstrates high ApEn, while a series containing repetitive patterns shows low ApEn.

The results of the ApEn estimation for the 400 experimental and shuffled trajectories are shown in Fig. 5 (*Supplementary Information*, Tables S6 and S7, respectively). In Fig. 5*A*, the ApEn values corresponding to intervals with fewer than 300 data points were omitted from the heatmaps. This exclusion is due to ApEn requiring a minimum of 10^m^ data points (where m=2 in our calculations) to yield meaningful results (Pincus et al., 1991). Notably, such short time series (with a length of ≤200 points) are considered unreliable for ApEn analysis (Yentes et al., 2013). The heatmaps illustrate the Approximate K-S entropy for the experimental (top row) and shuffled trajectories (bottom row) from cells (left panels) and cytoplasts (right panels), computed for 82 different time windows (intervals) with increasing length (we increased the interval duration by 25 seconds at each iteration). The intervals have ApEn values ranging from 10^−4^ (blue) to 0.46 (red) for experimental trajectories and from 0.12 (blue) to 2.13 (red) for shuffled trajectories. These results reveal that almost all the experimental series have very low entropy. To enhance visualization, we included interval ApEn values corresponding to the 300 first data points in the heatmap colormap ranges of Fig. 5*A*. The ApEn analysis of experimental trajectories yielded a narrow spectrum of low values ranging from 10^−4^ to 0.02, an overall median/IQR of 0.00/0.00 and a median/IQR of 0.00/0.00 for cells and 0.00/0.00 for cytoplasts.

**Fig. 5.**
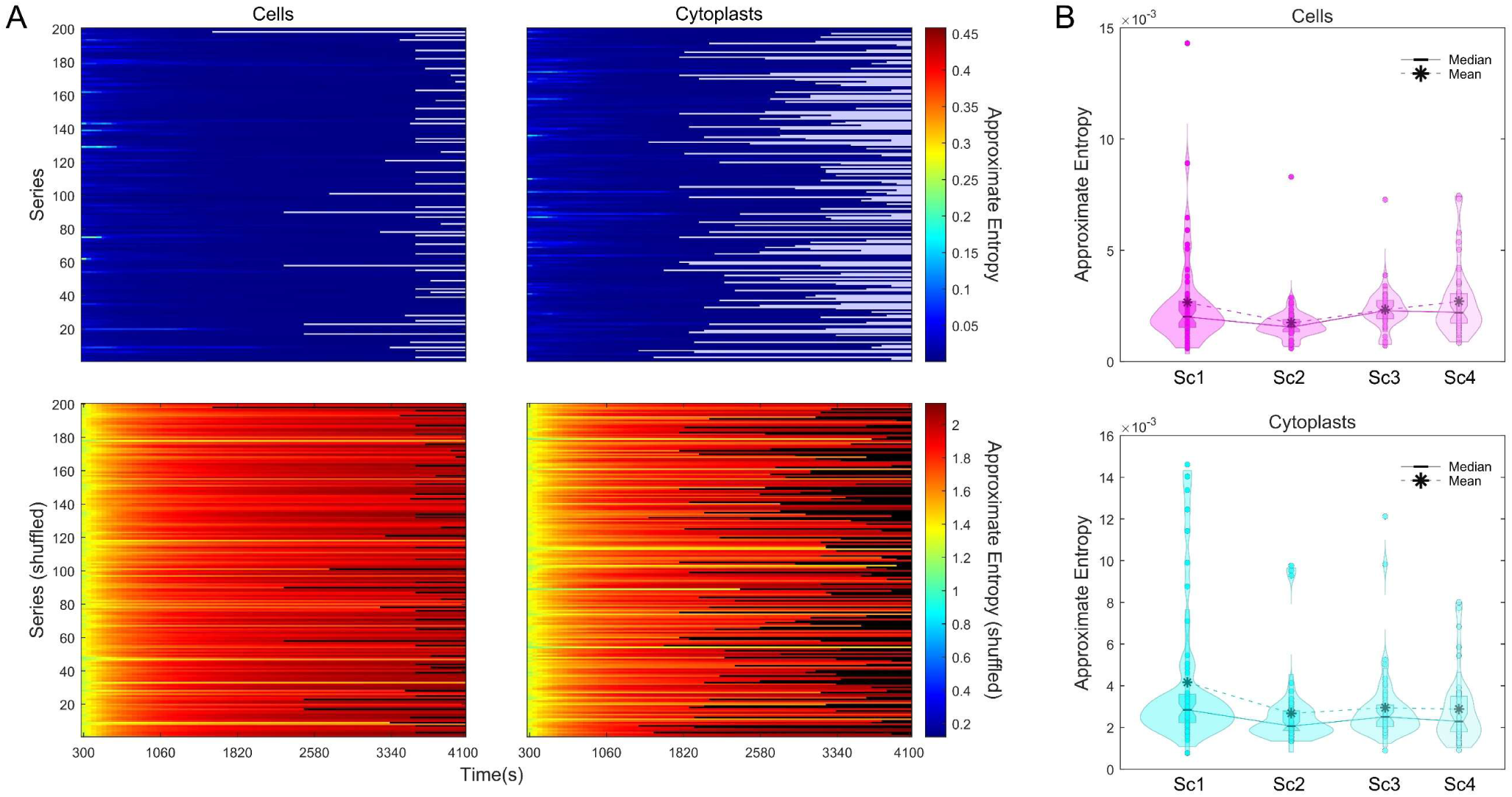
Unpredictability in cellular migration. *(A)* Heatmaps of Approximate Entropy values for all 400 experimental (upper row panels) and shuffled (lower row panels) cell (left panels) and cytoplast (right panels) trajectories. Approximate Entropy (ApEn) is a statistical tool used to determine the consistency and foreseeability of a time series. A signal with recurring patterns will have a low ApEn, indicating regularity, whereas a high ApEn is associated with an irregular and less predictable series. Every row in the heatmaps corresponds to one individual trajectory and the 82 columns display the number of data points used for Approximate Entropy calculation, increased in 50 data points (25 seconds) at every iteration. *(B)* Violin plots display the estimated distribution, mean and median of the Approximate Entropy values for all the experimental cell and cytoplast trajectories. “Sc1” Scenario One (no stimuli); “Sc2” Scenario Two (galvanotaxis under an electric field); “Sc3” Scenario Three (chemotaxis along a peptide gradient) and “Sc4” Scenario Four (simultaneous galvanotactic and chemotactic cues).

Conversely, the ApEn for shuffled trajectories had a range of very high values (from 1.30 to 2.13, median/IQR of 1.96/0.20) compared to the values for experimental trajectories (*Supplementary Information*, Table S7). Our analysis verifies the existence of a complex structure with high information content in the move-step sequences of the cell migration trajectories. Moreover, the results of a Wilcoxon test comparing the corresponding ApEn value distributions of the experimental and shuffled trajectories (p-value≅0 and Z=−24.48) evidence that the complex dynamic structure seen in the move-step series was extremely improbable to be random.

### 8. Persistence in cellular migration

Another key feature of the systemic cellular migration movements in unicellular organisms is the long-range memory effects or persistence (d’Alessandro et al., 2021; van Haastert, 2021). The Detrended Fluctuation Analysis (DFA) (see *Materials and Methods*) is a widely used method for detecting persistent effects in physiological time series.

For a specified observation scale, denoted as ℓ, DFA computes the function *F*(ℓ) to quantify the fluctuations of the time series around its local trend. If the time series exhibits scaling properties, then *F*(ℓ) ∼ ℓ^(y)^ asymptotically, where *γ* is the scaling exponent. This exponent is typically estimated as the slope of a linear fit in the log-log plot of *F*(ℓ) versus ℓ. Therefore, *γ* indicates the degree of long-term memory effects and describes the underlying dynamical system. Values near 0.5 imply the lack of long-range correlations, while when 1.5 < *γ* < 2, the process shows positive long-range persistence (Hardstone et al., 2012) (Fig. 6*A*). By applying this quantitative method, we detected the existence of long-range persistence in all experimental trajectories (*Supplementary Information*, Table S8), with an overall *γ* median/IQR value of 1.82/0.08. In particular, the median/IQR DFA scaling parameter *γ* was determined to be 1.82/0.08 for cells and 1.82/0.09 for cytoplasts (Fig. 6*B* and *Supplementary Information*, Table S8), thereby indicating that all the move-step series display trend-strengthening behavior.

**Fig. 6.**
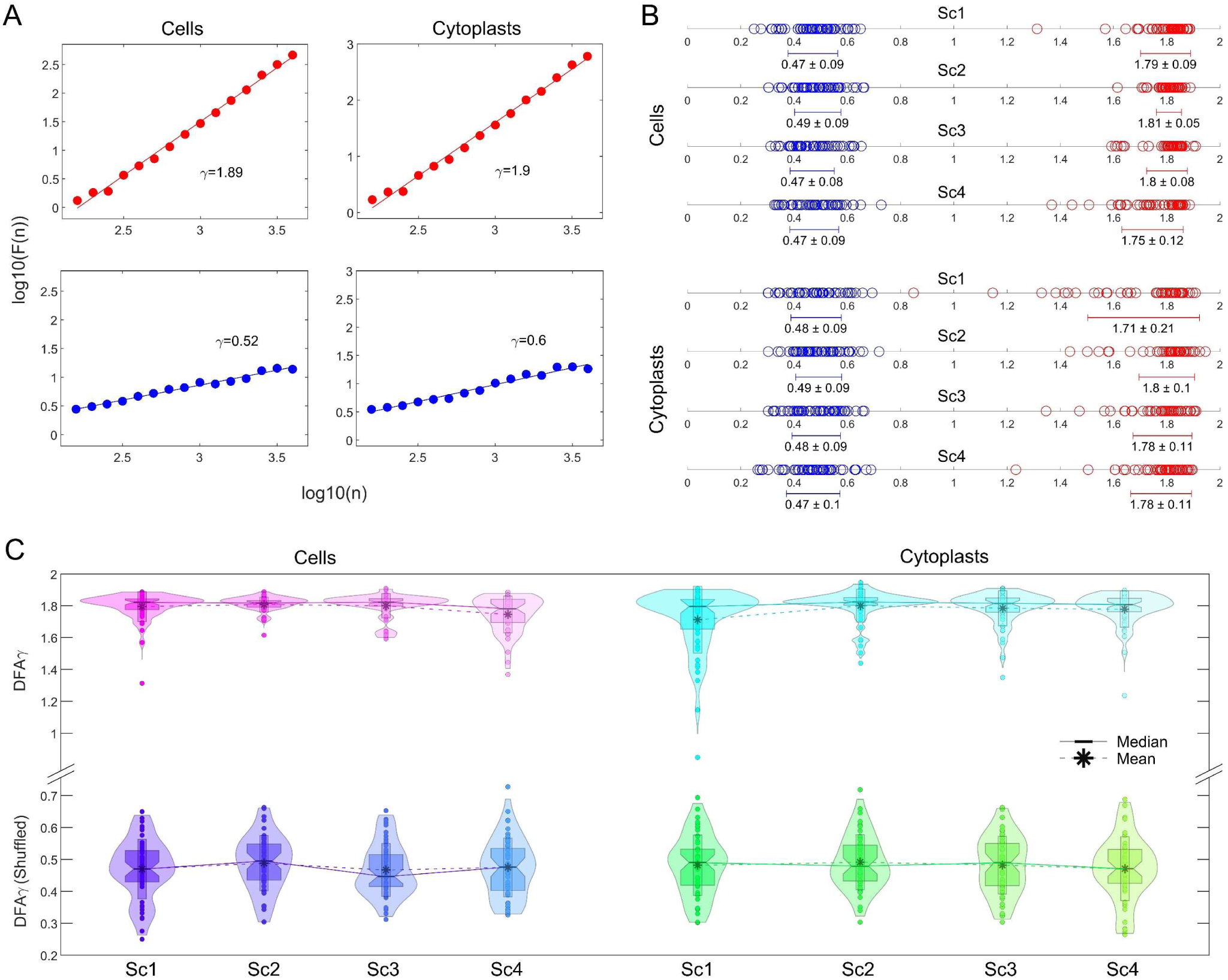
Long-term memory influences the migratory behavior of cells. *(A)* The detrended fluctuation parameter *F*(*n*) and the window size *n* for both a typical cell (left panels) and cytoplast (right panels) in the absence of stimuli (Sc1) are displayed on a log-log chart. The large scaling exponent *γ* values indicate long-range memory effects in both cells and cytoplasts. By shuffling, the original trajectories lost their memory information, which made the *γ* values decrease from *γ* = 1.89 to *γ* = 0.52 in the cell and from *γ* = 1.9 to *γ* = 0.6 in the cytoplast. *(B)* The scaling exponent *γ* value (and its average ± SD) for all cells and cytoplasts in every experimental scenario (Sc1*–*Sc4) is shown, experimental values in red and shuffled values in blue. *(C)* The violin plots depict the approximate distribution of scaling exponent *γ* values, along with their mean and median, for all experimental (top panel) and shuffled (lower panel) cell (left) and cytoplast (right) trajectories. “Sc1” Scenario One (no stimuli); “Sc2” Scenario Two (galvanotaxis under an electric field); “Sc3” Scenario Three (chemotaxis along a peptide gradient) and “Sc4” Scenario Four (simultaneous galvanotactic and chemotactic cues).

The strong correlation values observed in the experimental migration series disappeared after shuffling (see Fig. 6*B* and *C*, and *Supplementary Information*, Table S9 for more details), with *γ* overall median/IQR of 0.48/0.12. This finding verifies that the complex locomotion structure, characterized by well-structured move-step sequences and persistent dynamics observed in the migration trajectories of the two cell groups, is not due to a random chance (p-value≅0, Z=24.48).

### 9. Kinetic properties of cellular motion

We measured the Intensity of the response (IR), the directionality ratio (DR) and the average speed (AS) of amoebae (Fig. 7*A-C*) to evaluate some kinematic aspects of the cell migration trajectories.

**Fig. 7.**
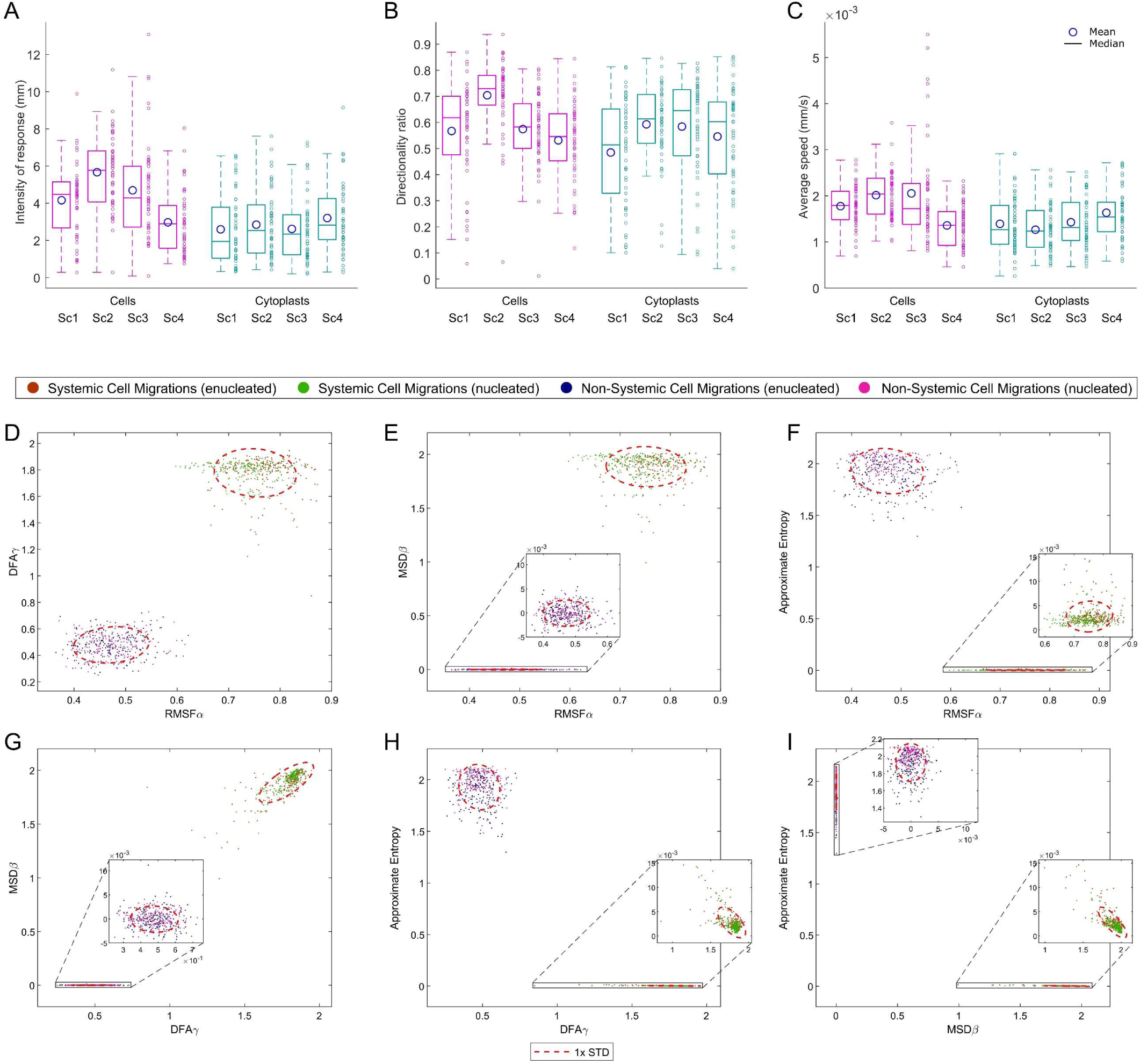
Kinematic characteristics and systemic properties of cell migration. *(A–C)* Group boxplots represent the distributions of all experimental values for three key cytokinetic metrics, which unveil fundamental information about the traits of cell migration—*(A)* Intensity of the response, *(B)* Directionality ratio, and *(C)* Average speed—for the two cell types (enucleated and non-enucleated) under all four experimental scenarios (Sc1, Sc2, Sc3, and Sc4). *(D–I)* The results of the analysis of the 400 experimental trajectories, with key metrics considered in our work, namely RMSF Alpha, DFA Gamma, MSD Beta, and Approximate Entropy, is juxtaposed with the corresponding shuffled data where the systemic informational framework was entirely disrupted. The values for cells and cytoplasts overlap in all comparisons, being qualitatively very similar. Ellipses (in red) representing one times the standard deviation of the data in each data “cloud” have been drawn to facilitate data interpretation. “Sc1” Scenario One (no stimuli); “Sc2” Scenario Two (galvanotaxis under an electric field); “Sc3” Scenario Three (chemotaxis along a peptide gradient) and “Sc4” Scenario Four (simultaneous galvanotactic and chemotactic cues).

The IR reflects the space covered by the cell, and specifically, we computed the modulus of the trajectories to indicate the response magnitude. The median/IQR IR was 4.31/3.04 for cells and 2.47/2.42 for cytoplasts. Then, we examined the DR, which calculates trajectory linearity values ranging between 0 (for completely curved trajectories) and 1 (for completely straight trajectories) by using the initial and final positions of the path taken by the amoeba. Experimental values ranged from 0.01 to 0.94 (median/IQR 0.62/0.21) for cells and 0.04 to 0.85 for cytoplasts (median/IQR 0.60/0.28). Lastly, we estimated the average speed (AS) of the trajectories, which varied between 10^−4^ and 0.01 mm/s (median/IQR 0.00/10^−4^) for cells and 10^−4^ and 0.00 mm/s (median/IQR 0.00/10^−4^) for cytoplasts.

### 10. Systemic properties of cell displacement

We compared the main metrics considered in our experimental study, such as RMSF Alpha, RMSF correlation time, DFA Gamma, MSD Beta, and Approximate Entropy, with data obtained in the corresponding shuffling procedures and represented them in Fig. 7*D*-*I*. The values for cells and cytoplasts overlap in all comparisons, showing high similarity. In each panel the shuffled and non-shuffled values for each metric could be clearly grouped and distinguished. This indicates that the shuffling process disorganized the inherent systemic information structure. The trajectories of the amoebae observed experimentally are totally distinct from those whose systemic properties were disrupted by the shuffling step (see Fig. 7*D-I*).

The results of all the quantitative analyses performed were qualitatively similar for both non-enucleated and enucleated cells (Table I).

**Table I.** Comparison between the migratory properties of non-enucleated and enucleated *A. proteus* cells.

| Quantitative tool | Non-enucleated (Median/IQR) | Enucleated (Median/IQR) |
| --- | --- | --- |
| RMSF $\alpha$ | 0.74/0.07 | 0.77/0.07 |
| Correlation time (in minutes) | 10.42/8.33 | 8.33/5.21 |
| MSD $\beta$ | 1.93/0.11 | 1.91/0.11 |
| DFA $\gamma$ | 1.82/0.08 | 1.82/0.09 |
| ApEn | 0.00/0.00 | 0.00/0.00 |
| Intensity of the Response (mm) | 4.31/3.04 | 2.47/2.42 |
| Directionality Ratio | 0.62/0.21 | 0.60/0.28 |
| Average Speed (mm/s) | 0.00/10<sup>-4</sup> | 0.00/10<sup>-4</sup> |

## Discussion

Migratory displacements represent a fundamental complex behavior of many cells being essential in Metazoan organisms. Deregulated locomotion movements are involved in important human diseases such as immunodeficiencies and cancer.

When unicellular organisms migrate, they must integrate a variety of external cues to carry out efficient movements, maximize displacement rate and develop optimal search trajectories to localize their targets. These strategies become particularly critical when the cells lack enough information on where the target is located. Despite the many research efforts made, many aspects of cellular migration remain largely unknown.

One of the key issues on cellular locomotion movements is whether this sophisticated behavior corresponds to a systemic property, and therefore is not found specifically in any of the functional biomolecular parts, partial cellular biomechanisms, or individual physiological processes. In this sense, recently, we have shown in a previous work that the migratory movements correspond to a complex emergent self-organized behavior at the cellular systemic level (De la Fuente et al., 2024).

Here, to continue delving into the study of the forces driving the locomotion movement of the cell, and to corroborate that the directional motility is regulated by integrative processes operating at a cellular global level, we have quantitatively analyzed the movement trajectories of 200 individual enucleated cells (cytoplasts) and 200 non-enucleated cells belonging to *Amoeba proteus* species (400 cells in total). All the locomotion trajectories have been obtained in four different experimental scenarios: in absence of stimuli, under a galvanotactic field, in chemotactic gradient, and under complex conditions such as simultaneous galvanotactic and chemotactic stimuli.

First, we have analyzed the long-range interdependence in the move-steps using the “rmsf” method rooted in Statistical Mechanics, drawing upon foundational concepts developed by Gibbs and Einstein (Gibbs, 1902; Einstein, 1909), that have been later applied in the analysis of different biological processes (Viswanathan et al., 1996; Ivanov et al., 1999, 2001). This method allowed us to quantify the long-range interdependence of amoebae move-steps. The obtained results have been similar in median/IQR for both non-enucleated cells (0.74/0.07) and cytoplasts (0.77/0.07). The analysis also revealed that each move-step in a trajectory is significantly influenced by its preceding move-step displacements, indicating robust dependencies over periods of 9.38/7.29 minutes (median/IQR, 10.42/8.33 minutes for non-enucleated cells and 8.33/5.21 for cytoplasts). Notably, across the four experimental scenarios considered all 400 unicellular organisms studied (enucleated and non-enucleated) exhibited non-trivial long-range correlations in their directional trajectories, a hallmark of an emergent systemic dynamics within the cell (van Haastert, 2021).

Furthermore, we identified anomalous behavior in migratory movements using the Mean Squared Displacement (MSD) method, also proposed by Einstein (Einstein, 1905). Displacement trajectories that do not result in a linear MSD can be considered as non-trivial, and specifically, the anomalous nature of cellular locomotion dynamics can be detected by super-diffusion. Particularly, we have observed that both non-enucleated and enucleated amoebas show similar values of super-diffusion with 1.93/0.11 (median/IQR) for non-enucleated cells and 1.91/0.11 for cytoplasts. Likewise, the MSD is a proxy for the surface area explored by each cell over time, and is a measure related to the overall migration efficiency, which corresponds to adecuate movements during exploration of the extracellular medium (Faustino et al., 2007; Viswanathan et al., 2008; Dieterich et al., 2022).

We have also quantified the regularity and unpredictability of fluctuations during migratory displacements using Approximate Entropy (ApEn) (Pincus et al., 1991; Delgado-Bonal and Marshak, 2019). The results show values of 0.00/0.00 for non-enucleated cells and 0.00/0.00 for cytoplasts, which reveal substantial information in all analyzed trajectories. These values confirm the existence of a highly complex structure of move-step sequences in the amoebas. The entropy is remarkably low in all the directional movements, suggesting that the migration patterns operate at a higher complexity level. This observed dynamic complexity in trajectories is highly improbable to occur by chance (practically, p-value ≅ 0).

In addition, we have verified the presence of long-term memory effects in cellular migratory movements. The DFA fluctuation analysis (Peng et al., 1994) indicates that the scaling exponent *γ* has a median/IQR value of 1.82/0.08 for non-enucleated cells and 1.82/0.09 for cytoplasts. These results imply that move-step trajectories exhibit trend-reinforcing memory, i.e., past directional movements showing increased move-step values are likely to be followed by future increasing trends, and vice versa. In essence, the evolution of cell system trajectories is greatly influenced by prior system movements over extended time periods. Therefore, these DFA results also validate the presence of strong long-range correlations in locomotion movements observed by the “rmsf” analysis.

Finally, we have analyzed three main kinematic properties of cellular motion. The results of the “Intensity of the cellular response” (IR) analysis displayed values of 4.31/3.04 for non-enucleated cells and 2.47/2.42 for cytoplasts (median/IQR). The “Directionality ratio” study (DR) showed 0.62/0.21 for non-enucleated cells and 0.60/0.28 for cytoplasts. The results of the “Average speed” (AS) were 0.00/10^−4^ for non-enucleated cells and 0.00/10^−4^ for cytoplasts.

The values obtained from the analysis of “rmsf”, “MSD”, “DFA”, “ApN”, “DR” and “AS” for both the non-enucleated and enucleated amoebas displayed very similar values. However, “IR” showed slightly different values. About this question, note that enucleated cells lack the nucleus, a structure with important mechanic and sensory functions in cell migration. In addition, enucleation is a notably aggressive process that damages the plasmatic membrane, and during which part of the cytoplasm is extracted together with the nucleus (see Fig. 1). Cells are very robust organisms, and the enucleation by micromanipulators might lead to some slight quantitative differences between the migratory characteristics of nucleated and enucleated cells, particularly affecting the intensity of the migratory response of cytoplasts.

The quantitative studies carried out using advanced non-linear physical-mathematical tools rooted in Statistical Physics, unequivocally show that in both the non-enucleated and enucleated amoebas emerges a very complex dynamic structure in the migratory movements of all analyzed cells. Such structure is characterized by highly organized move-step sequences with very low level of entropy and high information, non-trivial long-range interdependence in the move-steps, strong anomalous super-diffusion dynamics, long-term memory effects with trend-reinforcing behavior, and efficient movements to explore the extracellular medium. These essential characteristics of the locomotion movements are a consequence of the self-organized dynamics intrinsic to all unicellular organisms.

Cells are open systems that operate far from the thermodynamic equilibrium and exchange energy-matter with the environment (De la Fuente, 2014; Skene, 2015; Goldbeter, 2018). Under these conditions, the cellular physiology is characterized by exhibiting a great number of non-linear enzymatic interactions and irreversible enzymatic processes in the metabolic networks allowing the cell system to become spatially and temporally self-organized (Goldbeter, 2002; Halley and Winkler, 2008; Misteli, 2009; De la Fuente, 2010, 2015; Kondepudi et al., 2020; De la Fuente et al., 2021). There are different types of self-ordered biochemical dynamic structures in the cell, the most studied are temporal molecular oscillations (ultradian oscillations), molecular circadian rhythms (with oscillatory periods close to 24 h), and spatial traveling waves (coherent oscillation of metabolite concentrations that propagates progressively across the intracellular medium) (Goldbeter, 2002, 2018; De la Fuente et al., 2014, 2021).

Examples of ultradian oscillations involved in the regulation of cell migration include rhythms in the concentration of cytoskeletal actin (Cardamone et al., 2011), myosin dynamics (Balland et al., 2005; Schillers et al., 2010), rhythmic cAMP levels (Simao et al., 2021), and oscillations in intracellular Ca^2+^ (Rondé et al., 2000; Savage et al., 2019). Temporal oscillations with a period close to 24h have also been associated with cell migration. For instance, circadian rhythms shaping the concentration of adhesion molecules and of the chemokine CCL21 (Holtkamp et al., 2021), oscillations in intracellular pericentrin protein levels (Nakazato et al., 2023), and daily rhythms in the concentration of actin cytoskeleton regulatory molecules (Hoyle et al., 2017). Spatiotemporal waves examples include intracellular Ca^2+^ waves (Rondé et al., 2000; Takayama and Onami, 2016), actin waves (Vicker, 2002; Asano et al., 2008; Stankevicins et al., 2020), PIP3 waves (Asano et al., 2008), NAD(P)H waves (Kindzelskii et al., 1998; Kindzelskii and Petty, 2002), and kinase waves (Aoki et al., 2017; Aikin et al., 2020).

Long-term correlations have been observed in different physiological processes and biological mechanisms as, for instance, in the intracellular transport pathway (Ludington et al., 2013), NADPH levels (Ramanujan et al., 2006), glycolytic pathway (de la Fuente et al., 1998a; De la Fuente et al., 1998, 1999b; De la Fuente and Cortes, 2012), calcium activated potassium channels (Bandeira et al., 2008), and enzymatic dissipative networks (De la Fuente et al., 1999a, 2009), neural activity (de la Fuente et al., 2006), and complex biochemical dynamics (de la Fuente et al., 1998b). Furthermore, emergent systemic properties have been constated in cellular migratory displacements (De la Fuente et al., 2019a; Carrasco-Pujante et al., 2021).

On the other hand, the fact that cytoplasts preserved the dynamic properties of their migratory trajectories as the non-enucleated cells suggests that the nuclear genetic activity has a minor role in regulation of the locomotion displacements of cells. This result reinforces other previous studies in which similar migratory patterns were observed between cytoplasts belonging to different species and the corresponding non-enucleated cells (Graham et al., 2018; De la Fuente et al., 2019b).

In short, our experimental quantitative study indicates that migration is a complex self-regulated systemic property of cells, carefully regulated at a global level, which seems to depend on the cooperative non-linear interaction of most, if not all, molecular components of cells. This result does not invalidate the notable scientific importance of all studies performed on the influence of individual and particular molecular-physiological processes on cell migration.

Cell migration is a central issue in many human physiological and pathological processes, and we consider that new researches combining migratory systemic dynamics with molecular studies may be crucial for the development of next-generation, efficient cellular therapies for migration disorders.

## Materials and Methods

**Cell cultures.** Amoeba proteus (Carolina Biological Supply, #131306) were cultured at 21°C in Ø100 × 20 mm Petri dishes (Corning® CLS430167) containing Chalkley’s simplified medium (Korohoda et al., 2000) (NaCl, 1.4 mM; KCl, 0.026 mM; CaCl2, 0.01 mM) and one heat-treated wheat grain.

### Experimental set-up arrangement

A schematic representation of the experimental arrangement is presented in Figures S1 and S2. The setup involves two electrophoresis blocks (BIO-RAD Mini-Sub cell GT) put together in such a way that the first electrophoresis block is connected to a BIO-RAD Model 1000/500 power supply unit and to the second block by two ∼12-centimeter-long 2% agar bridges containing a 0.5 N KCl solution. Notably, the use of agar bridges and the removal of all electrodes from the second electrophoresis block serve to prevent direct contact between the moving cells and any metallic electrodes, thereby mitigating the risk of ionic contamination.

All experimental replicates were conducted using a maximum of eight cells and had a duration of 34 minutes and 10 seconds. Throughout this duration, cell behavior was meticulously recorded using a digital camera. Prior to commencing each experiment, amoebae were subjected to a 24-hour starvation period in fresh, non-nutritive simplified Chalkley’s medium. Only cells exhibiting optimal vitality (characterized by motility and spindle-shaped morphology) were selected for inclusion in the experiments. Notably, any deviations from optimal culture and experimental conditions, as well as mechanical anomalies within the recording system, led to the exclusion of approximately 10% of the experimental replicates. These invalidated replicates were not considered in the subsequent analysis.

The experiments were conducted within a custom-devised experimental glass chamber, put over the central elevated platform of the second electrophoresis block, where the amoebae were placed and could freely crawl. It consisted of a modified standard glass slide of dimensions 25 × 75 × 1 mm to which two 24 × 60 × 0.17 mm cover glasses were glued with silicone adhesive. After allowing the assembly to dry for 24 hours, the overhanging portions (measuring 20 × 60 × 0.17 mm) from both cover glasses were cut off, resulting in two 4 × 60 × 0.17 mm glass sheets that served as the chamber’s sidewalls. Finally, three additional smaller glass pieces, crafted by carefully trimming three 24 × 60 × 0.17 mm cover glasses, were placed over the modified glass slide: a central piece measuring 3 × 24 × 0.17 mm and two larger pieces flanking it, each measuring 24 × 40 × 0.17 mm. The two flanking pieces could be sled to close or open the experimental glass chamber to respectively facilitate the establishment of a laminar flow or allow for the placement and extraction of cells.

New agar bridges were employed in each iteration to prevent contamination and minimize conductivity loss. This precautionary measure aimed to avoid the infiltration of KCl from the simplified Chalkley’s medium (which possesses a significantly lower osmotic concentration than the agar bridges). Additionally, a dedicated electrophoresis block was exclusively allocated for experiments involving chemotaxis, while another block served specifically for galvanotaxis. Furthermore, the entire experimental setup, including both the electrophoresis blocks and the experimental glass chamber, underwent meticulous cleaning and reassembly after each experimental replication.

Prior to each experimental replicate, the modified glass slide was affixed to the upper surface of the central platform within the second electrophoresis block. This attachment was achieved using a droplet of olive oil, applied to prevent both the medium and electric current from permeating beneath the experimental glass chamber. Subsequently, the central glass piece, measuring 3 × 24 × 0.17 mm, was gently positioned atop the center of the modified glass slide. Following this, amoebae were carefully washed in fresh simplified Chalkley’s medium and then placed beneath the central glass piece within the chamber. During this step, it is crucial to complete the process within 15 seconds to prevent the amoebae from adhering to the inner surface of the plastic micropipette tip, which could potentially damage their cellular membranes. To complete the assembly, two lateral sliding glass pieces, each measuring 24 × 40 × 0.17 mm, were positioned flanking the central piece, slightly overhanging the wells of the electrophoresis block. Subsequently, each well was meticulously filled with 75 ml of clean simplified Chalkley’s medium. The lateral glass pieces were gently pressed down using a micropipette tip until they made contact with the medium, which spread beneath them due to surface tension. Finally, the two lateral glass pieces were longitudinally slid until they touched the central piece where the amoebae were situated. This action effectively closed the glass chamber, establishing a connection between the media in both wells.

### Enucleation

Amoebae underwent a double washing in Simplified Chalkley’s medium and were subsequently enucleated on 35 × 10 mm Petri dishes using a Sutter Instruments Co. MP-285 micromanipulator and an Olympus CK40 microscope. The enucleation procedure involved delicately introducing a slender glass pipette into the amoeba’s cytoplasm through the cell membrane, followed by manual extraction of the nucleus (See Fig. 1*C* and *D* for more information). It is important to acknowledge that the enucleation process is notably forceful, resulting in significant damage to the plasmatic membrane and the removal of a portion of the cytoplasm along with the nucleus. Consequently, this technique was limited to a maximum of two attempts per cell. Once successfully enucleated, the cytoplasts remained undisturbed for approximately 2 minutes.

### Fluorescent staining to confirm enucleation

To confirm nucleus removal, enucleated cells were stained with DAPI after each experiment and photographed in a fluorescence microscopy together with an also DAPI-stained control cell with its nucleus intact. Amoebae were first fixated in 4% paraformaldehyde (ENMA Bio Ltd., #28794.295) for 5 minutes. Following fixation, they were simultaneously permeabilized with Triton-X 100 (0.1%) and stained with DAPI (1 µg/ml) for 10 minutes. Finally, observation was carried out using an Olympus IX71 inverted fluorescence microscope at the High Resolution and Analytic Microscopy Service (SGIker) of the University of the Basque Country (UPV/EHU).

### Galvanotactic stimulus

A stable 60V direct current was applied throughout the experimental setup for the duration of the experiment. The galvanotactic stimulus can have traumatic effects on amoebal behavior. To mitigate these effects, we implemented four crucial steps in our experiments involving galvanotaxis (see *Supplementary Information*, Figure S1*A* for more detail):

1. Stable Power Supply: We programmed the power supply unit to deliver a consistent 60V direct current.
2. Chamber Customization: By adjusting the cross-sectional area of the experimental glass chamber through changing the amount of silicone adhesive used, we controlled the height of the longitudinal walls of the modified glass slide.
3. Resistance Management: We installed a variable 1 MΩ resistor and an ammeter in series (in that order). Real-time adjustments were made by manually turning the variable resistor’s screw to fine-tune the overall resistance of the setup. This ensured a stable 60V electric potential and optimal current intensity values of 70-74 µA throughout the galvanotactic experiments.
4. Immediate Termination: Once the recording concluded, the electric current was promptly stopped.

Interestingly, certain amoebae populations exhibited anomalous, inverted, or null responses to the galvanotactic stimulus. To exclude them from our study, we conducted a 5-minute galvanotactic test using intensity and voltage values within the optimal range before carrying out any experiments where amoebae were exposed to galvanotaxis for the first time.

### Chemotactic peptide gradient

To establish a chemotactic peptide gradient, we introduced 750 µl of nFMLP (Sigma-Aldrich, #F3506) at a concentration of 2 × 10⁻⁶ M into one well of the second electrophoresis block. The medium in this well was promptly stirred to ensure thorough mixing of the peptide until the amoebae exhibited a response.

To evaluate the nFMLP peptide gradient concentration, we conducted an experiment where 60 µL of medium was sampled at specific time intervals (0, 2, 5, 10, 15, 20, and 30 minutes) following the addition of nFMLP. These samples were extracted from the center of the experimental glass chamber through a narrow gap created by slightly shifting the lateral glass piece adjacent to the central glass component. By referencing known fluorescein-tagged peptide concentration values from a standard curve, we extrapolated the nFMLP concentration at each time point. The procedure involved taking two samples at each time interval and repeating the entire process three times, resulting in a total of six measurements for each time point. Fluorescence measurements were performed at 460/528 excitation/emission wavelengths using 96-well glass bottom black plates (Cellvis, #P96-1.5H-N) and a SynergyHTX plate reader (BIOTEK), following the established protocol by Green and Sambrook (Green and Sambrook, 2012).

### Trajectory recording and analysis

Each enucleated and non-enucleated amoeba was meticulously placed on an individual nutrient-free Petri dish. Subsequently, the motility of each cell was meticulously documented using a MU500 AmScope digital camera, which was affixed to the trinocular port of an AmScope SM-2T stereomicroscope. The stereoscope’s trinocular port includes a built-in 0.5 reduction lens, which added to a 0.75× reduction Barlow lens attached to the stereoscope’s objective allowed us to increase the field of view to approximately 3.3 × 2.5 cm. Data were collected at a rate of two images per second over a duration of 34 minutes and 10 seconds (4100 frames). The inclusion criterion stipulated that only trajectories exhibiting a minimum of 15 minutes (1800 frames) of active movement under the camera would be considered. To ensure precision, manual tracking was conducted using the TrackMate (Tinevez et al., 2017) software within ImageJ. This approach was preferred over automated tracking software, which is often prone to inaccuracies (Hilsenbeck et al., 2016), and to require manual error correction (Kok et al., 2020). Each tracked trajectory corresponded to a distinct amoeba, and no individual amoeba was recorded more than once. A total of 400 digitized trajectories, evenly divided into two groups (enucleated or cytoplasts and non-enucleated or cells), were subjected to analysis.

### Statistical analyses

In our study, we initially evaluated the normality of the distribution of our quantitative data using the Kolmogorov-Smirnov test for single samples. However, due to the rejection of normality, we proceeded to assess the significance of our quantitative findings using two non-parametric tests: The Kruskal-Wallis Test to compare multiple groups and the Wilcoxon Rank-Sum test for pairwise comparisons between groups. Since both tests are non-parametric, we reported our results in terms of median and interquartile range (IQR), rather than the traditional mean ± standard deviation. Additionally, we included p-values and Z statistics to provide a comprehensive understanding of the statistical significance.

## Data and code availability

This study did not generate any code.

Any additional information required to reanalyze the data reported in this paper is available from the lead contact upon request.

## Acknowledgments

We would like to thank Florentino Onandía Yague for valuable advice related to the experimental setup. This work was supported by grant US21/27 from the University of Basque Country (UPV/EHU) and Basque Center of Applied Mathematics. In addition, this work was supported by Basque Government funding, grant IT456-22.

## Author contributions

JC-P, and MF: performed the experiments; CB, MF and IMDF: designed the setup; CB methodology with experimental glass chamber; MF and JC-P: cell cultures and cellular behavior advice; JC-P: performed the digitalization of trajectories; IM, JMC, IMDF: designed quantitative analysis; JC-P, IM, JMC and BC-P: performed the quantitative studies; GPY, JIL, AP-S, JC-P, IM, JMC and IMDF: analysis and design of the research mapping; all authors wrote the manuscript and agreed with its submission; IMDF: conceived, designed and directed the investigation.

---

**Fig. S1.**
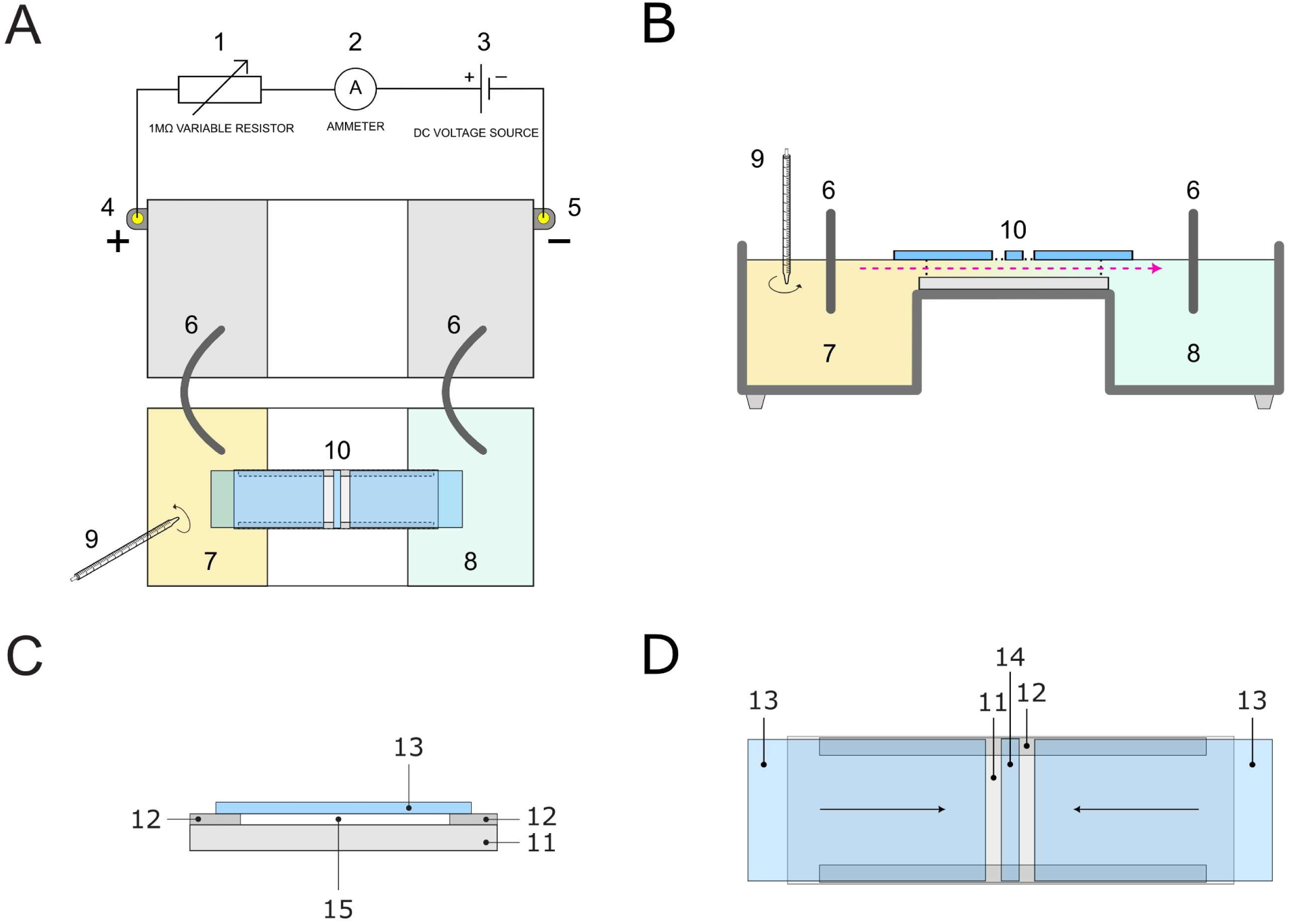
Experimental set-up configuration. (A) Top view of the experimental set-up. (B) Full section of the electrophoresis block where the amoebae were placed. 1: BIO-RAD model 1000/500 power supply unit, configured to provide a stable DC Voltage output; 2: Variable resistor (1 MΩ) to adjust the amount of circulating electric current (intensity); 3: ammeter to keep track of electric current values over time; 4: positive electrode (anode); 5: negative electrode (cathode); 6: agar + KCl bridges; 7: well were the peptide was added; 8: well containing fresh Chalkley’s medium, towards which the chemical gradient spreads over time; 9: a stirrer was utilized for the adequate blending of the chemotactic peptide in the medium; 10: experimental glass chamber, the direction and trajectory of the laminar flow is indicated by the pink arrow. (C) Sectional view of the experimental glass chamber. (D) Top view of the experimental glass chamber. 11: standard glass slide; 12: two longitudinal 4 × 60 × 0.17 mm glass stripes glued to *6* with silicone adhesive; 13: lateral sliding glass pieces; 14: central 3 × 24 × 0.17 mm glass stripe where the amoebae were placed underneath. 15: cross-sectional area of the experimental glass chamber. See *Materials and Methods* for more detail.

**Figure S2.**
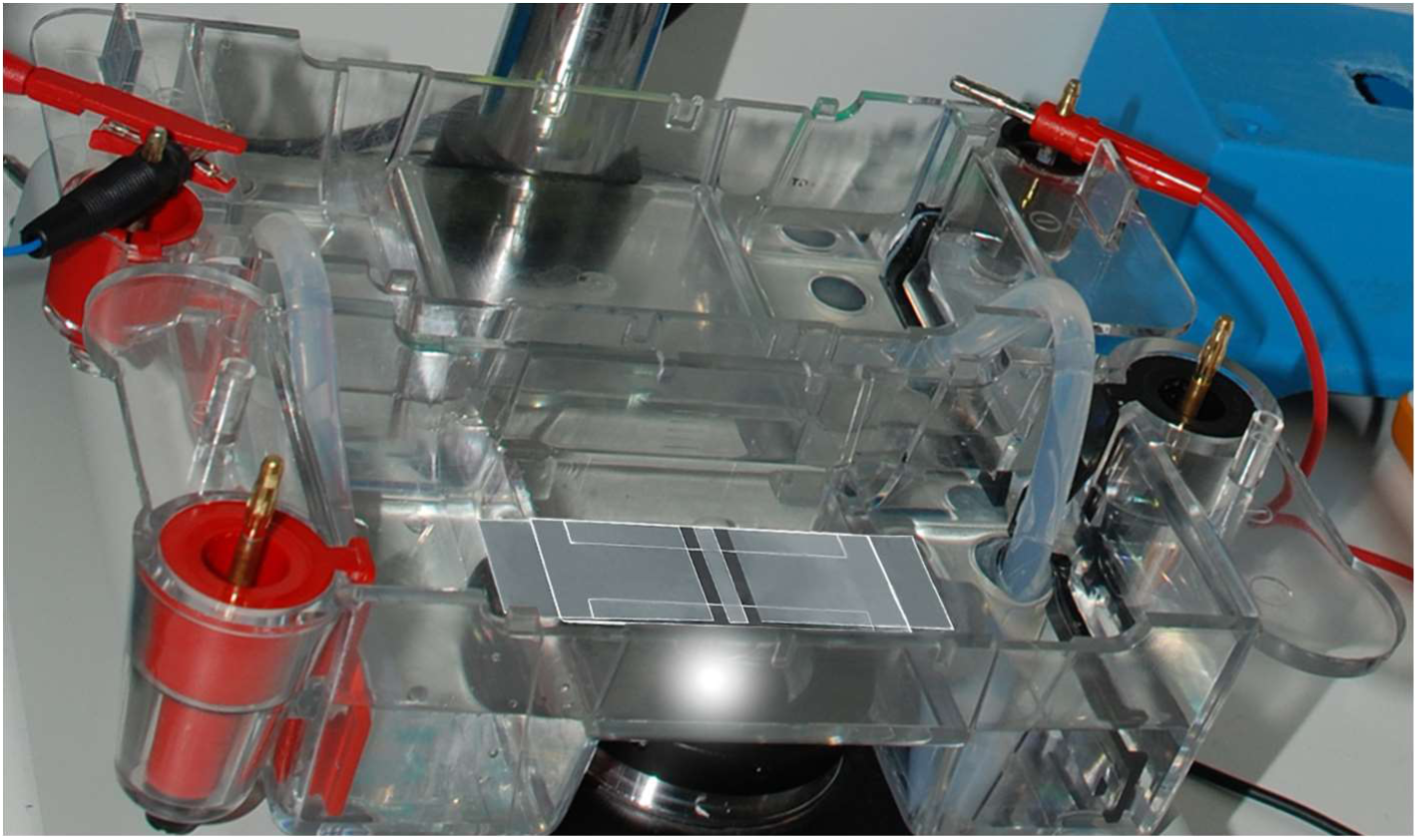
Experimental set-up under experimental conditions, related to Figure S1. Set-up layout and its components, illustrated in Figure S1, as seen under experimental conditions.

**Figure S3.**
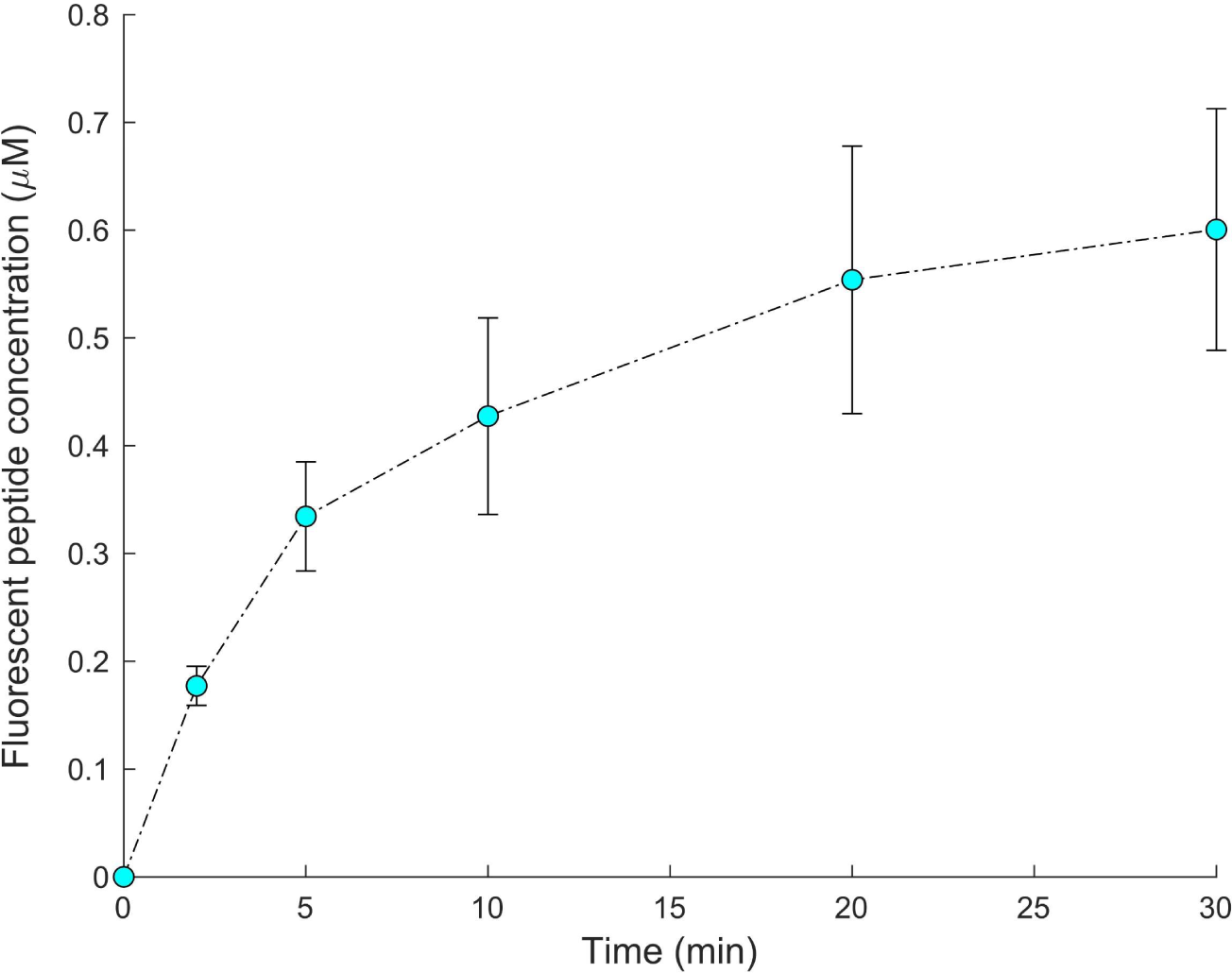
Fluorescein-tagged chemotactic peptide gradient evolution. We measured the temporal dynamics of peptide concentration at the center of the experimental glass chamber. Measurements were taken at specific time intervals (0, 2, 5, 10, 20, and 30 minutes) at the central location where amoebae were positioned. Each data point represents the average concentration (± standard deviation) obtained from six measurements (duplicate sampling in three separate experimental replicates). Notably, the peptide concentration exhibits an initial rise to approximately 0.2 μM within two minutes of establishing laminar flow. Subsequently, it further increases to 0.6 μM by the end of the 30-minute experiment.

**Table S1.** RMSF scaling exponent α values of the 400 experimental amoeba trajectories.

| # | Cells |  |  |  | Cytoplasts |  |  |  |
| --- | --- | --- | --- | --- | --- | --- | --- | --- |
|  | Sc1 | Sc2 | Sc3 | Sc4 | Sc1 | Sc2 | Sc3 | Sc4 |
| 1 | 0.742 | 0.725 | 0.652 | 0.769 | 0.672 | 0.667 | 0.757 | 0.838 |
| 2 | 0.781 | 0.761 | 0.75 | 0.738 | 0.766 | 0.819 | 0.78 | 0.776 |
| 3 | 0.824 | 0.737 | 0.742 | 0.798 | 0.666 | 0.726 | 0.774 | 0.781 |
| 4 | 0.777 | 0.749 | 0.76 | 0.709 | 0.827 | 0.772 | 0.714 | 0.772 |
| 5 | 0.822 | 0.79 | 0.743 | 0.805 | 0.764 | 0.688 | 0.732 | 0.781 |
| 6 | 0.868 | 0.739 | 0.751 | 0.737 | 0.813 | 0.757 | 0.682 | 0.811 |
| 7 | 0.678 | 0.777 | 0.795 | 0.679 | 0.785 | 0.79 | 0.72 | 0.768 |
| 8 | 0.703 | 0.652 | 0.628 | 0.754 | 0.751 | 0.758 | 0.745 | 0.751 |
| 9 | 0.606 | 0.775 | 0.725 | 0.696 | 0.69 | 0.693 | 0.765 | 0.763 |
| 10 | 0.853 | 0.633 | 0.736 | 0.757 | 0.772 | 0.741 | 0.795 | 0.696 |
| 11 | 0.651 | 0.765 | 0.685 | 0.744 | 0.781 | 0.818 | 0.797 | 0.815 |
| 12 | 0.73 | 0.742 | 0.702 | 0.797 | 0.738 | 0.784 | 0.743 | 0.795 |
| 13 | 0.805 | 0.711 | 0.739 | 0.733 | 0.737 | 0.85 | 0.87 | 0.794 |
| 14 | 0.661 | 0.735 | 0.636 | 0.701 | 0.86 | 0.826 | 0.786 | 0.66 |
| 15 | 0.676 | 0.671 | 0.777 | 0.783 | 0.733 | 0.804 | 0.754 | 0.698 |
| 16 | 0.69 | 0.718 | 0.727 | 0.8 | 0.818 | 0.744 | 0.734 | 0.848 |
| 17 | 0.753 | 0.69 | 0.763 | 0.659 | 0.744 | 0.711 | 0.75 | 0.796 |
| 18 | 0.725 | 0.661 | 0.759 | 0.768 | 0.817 | 0.849 | 0.813 | 0.714 |
| 19 | 0.738 | 0.744 | 0.597 | 0.666 | 0.754 | 0.729 | 0.806 | 0.797 |
| 20 | 0.664 | 0.659 | 0.693 | 0.83 | 0.86 | 0.795 | 0.851 | 0.776 |
| 21 | 0.8 | 0.691 | 0.748 | 0.69 | 0.714 | 0.767 | 0.775 | 0.817 |
| 22 | 0.801 | 0.624 | 0.782 | 0.713 | 0.803 | 0.835 | 0.823 | 0.75 |
| 23 | 0.719 | 0.697 | 0.688 | 0.678 | 0.732 | 0.802 | 0.759 | 0.741 |
| 24 | 0.81 | 0.716 | 0.767 | 0.7 | 0.802 | 0.788 | 0.758 | 0.807 |
| 25 | 0.728 | 0.641 | 0.698 | 0.832 | 0.778 | 0.741 | 0.838 | 0.705 |
| 26 | 0.784 | 0.766 | 0.79 | 0.712 | 0.737 | 0.785 | 0.831 | 0.756 |
| 27 | 0.725 | 0.744 | 0.785 | 0.762 | 0.688 | 0.8 | 0.738 | 0.772 |
| 28 | 0.751 | 0.747 | 0.767 | 0.823 | 0.742 | 0.78 | 0.822 | 0.733 |
| 29 | 0.756 | 0.667 | 0.702 | 0.722 | 0.814 | 0.769 | 0.816 | 0.694 |
| 30 | 0.753 | 0.684 | 0.65 | 0.715 | 0.785 | 0.775 | 0.849 | 0.815 |
| 31 | 0.704 | 0.751 | 0.719 | 0.753 | 0.827 | 0.774 | 0.761 | 0.731 |
| 32 | 0.651 | 0.713 | 0.616 | 0.702 | 0.774 | 0.664 | 0.731 | 0.737 |
| 33 | 0.771 | 0.841 | 0.71 | 0.713 | 0.766 | 0.81 | 0.814 | 0.792 |
| 34 | 0.799 | 0.626 | 0.742 | 0.762 | 0.663 | 0.714 | 0.766 | 0.776 |
| 35 | 0.807 | 0.827 | 0.696 | 0.793 | 0.778 | 0.754 | 0.705 | 0.723 |
| 36 | 0.78 | 0.617 | 0.763 | 0.745 | 0.755 | 0.733 | 0.695 | 0.76 |
| 37 | 0.737 | 0.709 | 0.695 | 0.769 | 0.826 | 0.778 | 0.746 | 0.719 |
| 38 | 0.769 | 0.738 | 0.756 | 0.756 | 0.832 | 0.787 | 0.861 | 0.735 |
| 39 | 0.792 | 0.765 | 0.73 | 0.704 | 0.806 | 0.768 | 0.742 | 0.766 |
| 40 | 0.709 | 0.73 | 0.766 | 0.739 | 0.764 | 0.755 | 0.733 | 0.786 |
| 41 | 0.679 | 0.662 | 0.779 | 0.751 | 0.714 | 0.819 | 0.84 | 0.694 |
| 42 | 0.721 | 0.68 | 0.71 | 0.776 | 0.801 | 0.788 | 0.742 | 0.745 |
| 43 | 0.775 | 0.748 | 0.729 | 0.727 | 0.83 | 0.78 | 0.794 | 0.771 |
| 44 | 0.828 | 0.759 | 0.749 | 0.742 | 0.74 | 0.767 | 0.717 | 0.686 |
| 45 | 0.873 | 0.721 | 0.741 | 0.716 | 0.797 | 0.791 | 0.866 | 0.807 |
| 46 | 0.725 | 0.747 | 0.723 | 0.744 | 0.806 | 0.767 | 0.701 | 0.756 |
| 47 | 0.853 | 0.738 | 0.7 | 0.733 | 0.827 | 0.709 | 0.765 | 0.671 |
| 48 | 0.851 | 0.669 | 0.726 | 0.755 | 0.719 | 0.81 | 0.792 | 0.811 |
| 49 | 0.684 | 0.673 | 0.741 | 0.743 | 0.782 | 0.78 | 0.747 | 0.758 |
| 50 | 0.68 | 0.732 | 0.775 | 0.796 | 0.843 | 0.746 | 0.736 | 0.868 |

**Table S2.** RMSF scaling exponent α values of the 400 shuffled amoeba trajectories.

| # | Cells |  |  |  | Cytoplasts |  |  |  |
| --- | --- | --- | --- | --- | --- | --- | --- | --- |
|  | Sc1 | Sc2 | Sc3 | Sc4 | Sc1 | Sc2 | Sc3 | Sc4 |
| 1 | 0.44 | 0.51 | 0.435 | 0.449 | 0.536 | 0.431 | 0.41 | 0.484 |
| 2 | 0.428 | 0.48 | 0.537 | 0.517 | 0.394 | 0.515 | 0.365 | 0.431 |
| 3 | 0.445 | 0.493 | 0.443 | 0.512 | 0.411 | 0.524 | 0.414 | 0.513 |
| 4 | 0.415 | 0.518 | 0.455 | 0.574 | 0.497 | 0.404 | 0.466 | 0.553 |
| 5 | 0.529 | 0.403 | 0.497 | 0.446 | 0.481 | 0.517 | 0.543 | 0.461 |
| 6 | 0.462 | 0.43 | 0.503 | 0.469 | 0.473 | 0.433 | 0.468 | 0.505 |
| 7 | 0.405 | 0.402 | 0.44 | 0.429 | 0.396 | 0.46 | 0.387 | 0.445 |
| 8 | 0.443 | 0.46 | 0.379 | 0.474 | 0.456 | 0.535 | 0.414 | 0.525 |
| 9 | 0.482 | 0.561 | 0.401 | 0.512 | 0.527 | 0.47 | 0.622 | 0.514 |
| 10 | 0.548 | 0.535 | 0.52 | 0.487 | 0.448 | 0.475 | 0.507 | 0.521 |
| 11 | 0.553 | 0.483 | 0.496 | 0.615 | 0.533 | 0.405 | 0.439 | 0.454 |
| 12 | 0.468 | 0.408 | 0.467 | 0.52 | 0.521 | 0.469 | 0.506 | 0.536 |
| 13 | 0.464 | 0.451 | 0.447 | 0.498 | 0.437 | 0.474 | 0.55 | 0.524 |
| 14 | 0.502 | 0.444 | 0.583 | 0.447 | 0.532 | 0.509 | 0.433 | 0.43 |
| 15 | 0.422 | 0.446 | 0.563 | 0.405 | 0.569 | 0.416 | 0.487 | 0.54 |
| 16 | 0.477 | 0.496 | 0.505 | 0.53 | 0.561 | 0.467 | 0.549 | 0.471 |
| 17 | 0.427 | 0.454 | 0.465 | 0.459 | 0.481 | 0.421 | 0.438 | 0.504 |
| 18 | 0.481 | 0.503 | 0.493 | 0.49 | 0.405 | 0.506 | 0.502 | 0.402 |
| 19 | 0.512 | 0.471 | 0.497 | 0.523 | 0.44 | 0.387 | 0.474 | 0.459 |
| 20 | 0.521 | 0.406 | 0.438 | 0.436 | 0.488 | 0.48 | 0.425 | 0.418 |
| 21 | 0.4 | 0.426 | 0.405 | 0.558 | 0.373 | 0.454 | 0.533 | 0.47 |
| 22 | 0.385 | 0.405 | 0.504 | 0.462 | 0.436 | 0.484 | 0.534 | 0.401 |
| 23 | 0.436 | 0.471 | 0.47 | 0.466 | 0.515 | 0.502 | 0.509 | 0.573 |
| 24 | 0.431 | 0.485 | 0.505 | 0.434 | 0.495 | 0.497 | 0.524 | 0.504 |
| 25 | 0.492 | 0.477 | 0.461 | 0.442 | 0.503 | 0.442 | 0.426 | 0.45 |
| 26 | 0.587 | 0.377 | 0.429 | 0.456 | 0.456 | 0.409 | 0.449 | 0.503 |
| 27 | 0.39 | 0.441 | 0.447 | 0.612 | 0.529 | 0.469 | 0.469 | 0.468 |
| 28 | 0.447 | 0.498 | 0.488 | 0.5 | 0.437 | 0.432 | 0.558 | 0.451 |
| 29 | 0.473 | 0.504 | 0.446 | 0.42 | 0.5 | 0.488 | 0.428 | 0.482 |
| 30 | 0.522 | 0.449 | 0.517 | 0.594 | 0.609 | 0.486 | 0.425 | 0.44 |
| 31 | 0.471 | 0.423 | 0.498 | 0.41 | 0.457 | 0.514 | 0.499 | 0.431 |
| 32 | 0.448 | 0.489 | 0.503 | 0.532 | 0.477 | 0.398 | 0.466 | 0.495 |
| 33 | 0.517 | 0.416 | 0.408 | 0.429 | 0.519 | 0.404 | 0.598 | 0.442 |
| 34 | 0.491 | 0.395 | 0.508 | 0.48 | 0.476 | 0.48 | 0.431 | 0.538 |
| 35 | 0.529 | 0.421 | 0.519 | 0.545 | 0.395 | 0.53 | 0.457 | 0.495 |
| 36 | 0.429 | 0.454 | 0.54 | 0.484 | 0.397 | 0.531 | 0.58 | 0.522 |
| 37 | 0.494 | 0.402 | 0.449 | 0.473 | 0.411 | 0.487 | 0.444 | 0.469 |
| 38 | 0.484 | 0.458 | 0.468 | 0.457 | 0.48 | 0.567 | 0.548 | 0.486 |
| 39 | 0.47 | 0.463 | 0.456 | 0.484 | 0.396 | 0.421 | 0.453 | 0.508 |
| 40 | 0.468 | 0.447 | 0.443 | 0.41 | 0.486 | 0.505 | 0.412 | 0.453 |
| 41 | 0.47 | 0.4 | 0.466 | 0.495 | 0.453 | 0.521 | 0.449 | 0.45 |
| 42 | 0.512 | 0.51 | 0.468 | 0.427 | 0.422 | 0.431 | 0.456 | 0.406 |
| 43 | 0.475 | 0.599 | 0.403 | 0.475 | 0.472 | 0.398 | 0.503 | 0.508 |
| 44 | 0.512 | 0.415 | 0.48 | 0.385 | 0.468 | 0.491 | 0.455 | 0.407 |
| 45 | 0.486 | 0.371 | 0.485 | 0.541 | 0.439 | 0.515 | 0.543 | 0.425 |
| 46 | 0.444 | 0.477 | 0.444 | 0.449 | 0.47 | 0.521 | 0.443 | 0.546 |
| 47 | 0.48 | 0.421 | 0.454 | 0.495 | 0.433 | 0.423 | 0.428 | 0.51 |
| 48 | 0.539 | 0.45 | 0.444 | 0.487 | 0.47 | 0.451 | 0.464 | 0.401 |
| 49 | 0.538 | 0.509 | 0.436 | 0.408 | 0.451 | 0.491 | 0.457 | 0.445 |
| 50 | 0.467 | 0.425 | 0.455 | 0.438 | 0.503 | 0.466 | 0.431 | 0.407 |

**Table S3.** Long-range correlation span (memory time) in minutes for the 400 experimental amoeba trajectories.

| # | Cells |  |  |  | Cytoplasts |  |  |  |
| --- | --- | --- | --- | --- | --- | --- | --- | --- |
|  | Sc1 | Sc2 | Sc3 | Sc4 | Sc1 | Sc2 | Sc3 | Sc4 |
| 1 | 15.625 | 4.167 | 6.25 | 8.333 | 9.375 | 8.333 | 4.167 | 3.125 |
| 2 | 5.208 | 7.292 | 9.375 | 7.292 | 9.375 | 16.667 | 4.167 | 4.167 |
| 3 | 13.542 | 3.125 | 3.125 | 3.125 | 14.583 | 16.667 | 9.375 | 3.125 |
| 4 | 14.583 | 15.625 | 7.292 | 9.375 | 2.083 | 11.458 | 9.375 | 9.375 |
| 5 | 11.458 | 6.25 | 15.625 | 7.292 | 8.333 | 6.25 | 12.5 | 3.125 |
| 6 | 16.667 | 13.542 | 12.5 | 17.708 | 16.667 | 11.458 | 6.25 | 5.208 |
| 7 | 5.208 | 16.667 | 5.208 | 10.417 | 7.292 | 10.417 | 15.625 | 8.333 |
| 8 | 6.25 | 5.208 | 15.625 | 15.625 | 6.25 | 13.542 | 13.542 | 8.333 |
| 9 | 16.667 | 17.708 | 6.25 | 17.708 | 4.167 | 9.375 | 16.667 | 4.167 |
| 10 | 16.667 | 13.542 | 11.458 | 13.542 | 5.208 | 12.5 | 8.333 | 5.208 |
| 11 | 5.208 | 3.125 | 13.542 | 11.458 | 4.167 | 8.333 | 12.5 | 9.375 |
| 12 | 5.208 | 3.125 | 12.5 | 8.333 | 8.333 | 6.25 | 15.625 | 9.375 |
| 13 | 4.167 | 2.083 | 17.708 | 12.5 | 4.167 | 13.542 | 8.333 | 4.167 |
| 14 | 12.5 | 9.375 | 5.208 | 13.542 | 8.333 | 9.375 | 13.542 | 12.5 |
| 15 | 7.292 | 2.083 | 14.583 | 9.375 | 5.208 | 13.542 | 7.292 | 10.417 |
| 16 | 4.167 | 10.417 | 7.292 | 6.25 | 8.333 | 5.208 | 3.125 | 7.292 |
| 17 | 3.125 | 5.208 | 4.167 | 10.417 | 13.542 | 15.625 | 7.292 | 6.25 |
| 18 | 4.167 | 15.625 | 11.458 | 12.5 | 13.542 | 16.667 | 16.667 | 14.583 |
| 19 | 15.625 | 2.083 | 17.708 | 2.083 | 5.208 | 8.333 | 9.375 | 4.167 |
| 20 | 12.5 | 4.167 | 10.417 | 13.542 | 5.208 | 7.292 | 11.458 | 8.333 |
| 21 | 13.542 | 5.208 | 9.375 | 13.542 | 8.333 | 14.583 | 5.208 | 6.25 |
| 22 | 13.542 | 6.25 | 8.333 | 8.333 | 2.083 | 6.25 | 13.542 | 15.625 |
| 23 | 12.5 | 15.625 | 7.292 | 10.417 | 8.333 | 7.292 | 6.25 | 10.417 |
| 24 | 15.625 | 13.542 | 7.292 | 14.583 | 15.625 | 7.292 | 7.292 | 15.625 |
| 25 | 8.333 | 15.625 | 4.167 | 15.625 | 7.292 | 10.417 | 6.25 | 12.5 |
| 26 | 16.667 | 15.625 | 6.25 | 7.292 | 13.542 | 5.208 | 6.25 | 6.25 |
| 27 | 13.542 | 6.25 | 10.417 | 17.708 | 17.708 | 6.25 | 9.375 | 6.25 |
| 28 | 8.333 | 7.292 | 6.25 | 10.417 | 3.125 | 9.375 | 9.375 | 5.208 |
| 29 | 4.167 | 12.5 | 3.125 | 6.25 | 10.417 | 9.375 | 12.5 | 8.333 |
| 30 | 7.292 | 7.292 | 18.75 | 13.542 | 13.542 | 15.625 | 12.5 | 12.5 |
| 31 | 10.417 | 7.292 | 4.167 | 11.458 | 12.5 | 5.208 | 11.458 | 12.5 |
| 32 | 17.708 | 11.458 | 10.417 | 16.667 | 14.583 | 10.417 | 5.208 | 7.292 |
| 33 | 15.625 | 17.708 | 13.542 | 16.667 | 5.208 | 16.667 | 7.292 | 3.125 |
| 34 | 8.333 | 16.667 | 9.375 | 7.292 | 10.417 | 7.292 | 10.417 | 7.292 |
| 35 | 17.708 | 16.667 | 10.417 | 18.75 | 9.375 | 11.458 | 8.333 | 14.583 |
| 36 | 7.292 | 17.708 | 15.625 | 8.333 | 9.375 | 6.25 | 6.25 | 5.208 |
| 37 | 6.25 | 15.625 | 11.458 | 13.542 | 3.125 | 8.333 | 10.417 | 6.25 |
| 38 | 15.625 | 11.458 | 3.125 | 10.417 | 7.292 | 8.333 | 4.167 | 8.333 |
| 39 | 15.625 | 8.333 | 10.417 | 8.333 | 3.125 | 8.333 | 10.417 | 6.25 |
| 40 | 13.542 | 14.583 | 15.625 | 11.458 | 8.333 | 8.333 | 10.417 | 10.417 |
| 41 | 7.292 | 6.25 | 9.375 | 16.667 | 4.167 | 6.25 | 4.167 | 5.208 |
| 42 | 14.583 | 12.5 | 9.375 | 4.167 | 14.583 | 9.375 | 7.292 | 8.333 |
| 43 | 3.125 | 14.583 | 4.167 | 3.125 | 7.292 | 8.333 | 3.125 | 5.208 |
| 44 | 7.292 | 17.708 | 12.5 | 8.333 | 11.458 | 6.25 | 6.25 | 12.5 |
| 45 | 3.125 | 11.458 | 11.458 | 5.208 | 10.417 | 6.25 | 4.167 | 15.625 |
| 46 | 6.25 | 16.667 | 6.25 | 16.667 | 17.708 | 16.667 | 9.375 | 14.583 |
| 47 | 17.708 | 16.667 | 8.333 | 14.583 | 15.625 | 17.708 | 5.208 | 13.542 |
| 48 | 17.708 | 14.583 | 8.333 | 6.25 | 6.25 | 5.208 | 4.167 | 8.333 |
| 49 | 9.375 | 13.542 | 11.458 | 4.167 | 10.417 | 11.458 | 5.208 | 10.417 |
| 50 | 13.542 | 12.5 | 10.417 | 14.583 | 4.167 | 9.375 | 8.333 | 7.292 |

**Table S4.** MSD diffusion exponent β values of the 400 experimental amoeba trajectories.

| # | Cells |  |  |  | Cytoplasts |  |  |  |
| --- | --- | --- | --- | --- | --- | --- | --- | --- |
|  | Sc1 | Sc2 | Sc3 | Sc4 | Sc1 | Sc2 | Sc3 | Sc4 |
| 1 | 1.96 | 1.955 | 1.994 | 1.775 | 1.519 | 1.742 | 1.928 | 2.013 |
| 2 | 1.981 | 1.921 | 1.998 | 1.956 | 1.271 | 1.993 | 1.858 | 1.995 |
| 3 | 1.972 | 1.934 | 1.719 | 1.902 | 1.333 | 1.975 | 1.947 | 1.908 |
| 4 | 1.936 | 1.95 | 1.974 | 1.622 | 1.527 | 1.97 | 1.86 | 1.968 |
| 5 | 1.971 | 1.776 | 1.931 | 1.796 | 1.854 | 1.621 | 1.844 | 2.004 |
| 6 | 1.929 | 1.982 | 1.834 | 1.801 | 1.89 | 1.911 | 1.996 | 1.935 |
| 7 | 2.001 | 1.923 | 1.893 | 1.851 | 1.715 | 1.932 | 1.938 | 1.989 |
| 8 | 1.971 | 1.887 | 1.979 | 1.934 | 0.993 | 1.815 | 1.892 | 1.976 |
| 9 | 1.97 | 1.842 | 1.934 | 1.826 | 1.894 | 1.882 | 1.908 | 1.923 |
| 10 | 1.948 | 1.987 | 1.867 | 1.902 | 1.955 | 1.774 | 1.9 | 1.944 |
| 11 | 1.993 | 1.984 | 1.952 | 1.951 | 1.991 | 1.998 | 1.804 | 1.925 |
| 12 | 1.945 | 1.958 | 1.956 | 1.855 | 1.847 | 1.891 | 1.279 | 1.92 |
| 13 | 1.957 | 1.925 | 1.973 | 1.91 | 1.442 | 1.89 | 1.973 | 1.614 |
| 14 | 2.01 | 1.933 | 1.957 | 1.984 | 1.842 | 1.921 | 1.803 | 1.787 |
| 15 | 1.899 | 1.984 | 1.82 | 1.979 | 1.97 | 1.991 | 1.91 | 1.759 |
| 16 | 1.938 | 1.878 | 1.615 | 1.965 | 1.944 | 1.947 | 2.001 | 1.876 |
| 17 | 1.706 | 1.938 | 1.757 | 1.985 | 1.914 | 1.955 | 1.988 | 1.841 |
| 18 | 1.604 | 1.811 | 1.909 | 1.968 | 1.97 | 1.922 | 1.944 | 1.869 |
| 19 | 1.813 | 1.877 | 1.936 | 1.996 | 1.974 | 1.882 | 1.862 | 1.914 |
| 20 | 1.613 | 1.992 | 1.957 | 1.995 | 1.933 | 1.888 | 2.009 | 1.818 |
| 21 | 1.931 | 1.899 | 1.826 | 1.963 | 1.997 | 1.911 | 1.818 | 1.807 |
| 22 | 1.879 | 1.968 | 1.962 | 1.992 | 2.001 | 1.922 | 1.887 | 1.809 |
| 23 | 1.81 | 1.949 | 1.95 | 1.909 | 1.622 | 1.83 | 1.81 | 1.818 |
| 24 | 1.928 | 1.971 | 1.791 | 1.853 | 1.697 | 1.935 | 1.982 | 1.623 |
| 25 | 1.926 | 1.983 | 1.934 | 1.834 | 1.757 | 1.922 | 2.004 | 1.972 |
| 26 | 2.006 | 1.973 | 1.888 | 1.893 | 1.859 | 1.841 | 1.898 | 1.812 |
| 27 | 1.788 | 1.955 | 1.806 | 1.965 | 1.988 | 1.808 | 1.881 | 1.903 |
| 28 | 1.949 | 1.928 | 1.94 | 1.828 | 1.684 | 1.94 | 1.911 | 1.813 |
| 29 | 1.799 | 1.955 | 1.922 | 1.809 | 1.86 | 1.803 | 1.917 | 1.938 |
| 30 | 1.414 | 1.915 | 1.9 | 1.851 | 1.919 | 1.891 | 1.921 | 1.875 |
| 31 | 1.978 | 1.904 | 1.935 | 1.786 | 1.754 | 1.814 | 1.982 | 1.738 |
| 32 | 1.955 | 1.955 | 1.865 | 1.833 | 1.957 | 1.974 | 1.903 | 1.93 |
| 33 | 2.003 | 1.937 | 1.903 | 1.954 | 1.82 | 1.891 | 1.989 | 1.926 |
| 34 | 1.93 | 1.944 | 1.876 | 1.679 | 1.938 | 1.89 | 1.911 | 1.96 |
| 35 | 1.971 | 1.916 | 1.927 | 1.901 | 1.935 | 1.89 | 1.909 | 1.726 |
| 36 | 1.92 | 1.992 | 1.926 | 1.612 | 1.748 | 1.94 | 1.955 | 1.858 |
| 37 | 1.958 | 1.971 | 1.841 | 1.841 | 1.684 | 1.78 | 1.622 | 1.939 |
| 38 | 1.947 | 1.989 | 1.973 | 1.862 | 1.844 | 1.955 | 1.858 | 1.98 |
| 39 | 1.832 | 2.011 | 1.788 | 1.888 | 1.898 | 1.853 | 1.909 | 1.788 |
| 40 | 1.978 | 1.981 | 1.943 | 1.836 | 1.976 | 1.988 | 1.967 | 1.921 |
| 41 | 1.893 | 1.99 | 1.952 | 1.743 | 1.965 | 1.918 | 1.942 | 1.931 |
| 42 | 1.811 | 1.994 | 1.941 | 1.777 | 1.96 | 1.926 | 1.937 | 2.007 |
| 43 | 1.965 | 1.931 | 1.963 | 1.769 | 1.721 | 1.974 | 1.949 | 1.908 |
| 44 | 1.769 | 1.689 | 1.983 | 1.81 | 1.932 | 1.986 | 2.007 | 1.861 |
| 45 | 1.799 | 1.969 | 2.01 | 1.891 | 1.934 | 1.948 | 1.884 | 1.877 |
| 46 | 1.891 | 1.951 | 1.938 | 1.828 | 1.948 | 1.966 | 1.916 | 1.9 |
| 47 | 1.938 | 1.973 | 1.911 | 1.786 | 1.948 | 1.897 | 1.892 | 1.826 |
| 48 | 1.924 | 1.977 | 1.987 | 1.874 | 1.906 | 1.895 | 2.008 | 1.718 |
| 49 | 1.872 | 1.982 | 1.978 | 1.416 | 1.897 | 1.532 | 1.949 | 1.375 |
| 50 | 2.006 | 1.989 | 1.913 | 1.894 | 1.735 | 1.879 | 1.922 | 1.879 |

**Table S5.**
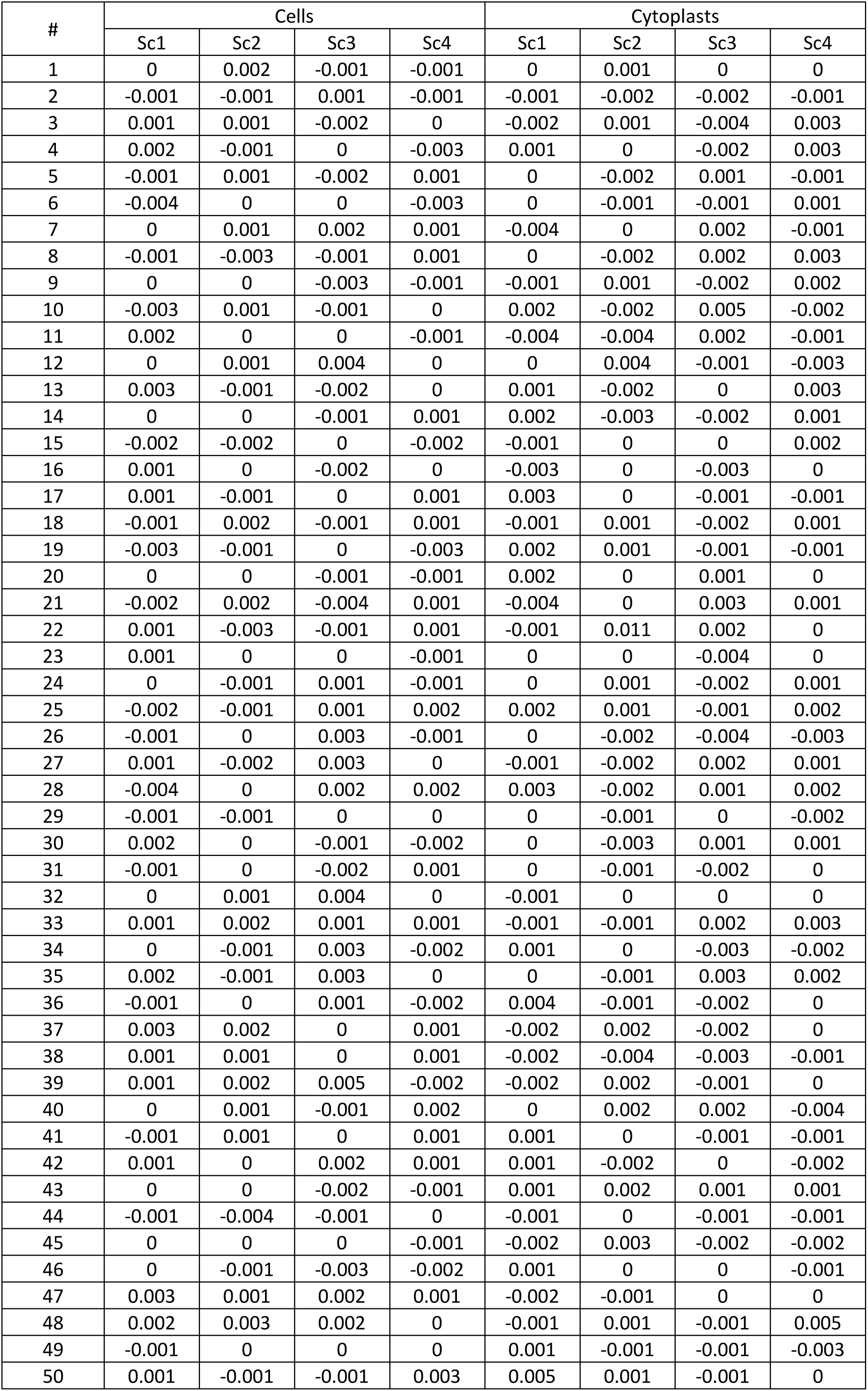
MSD diffusion exponent β values of the 400 shuffled amoeba trajectories.

**Table S6.**
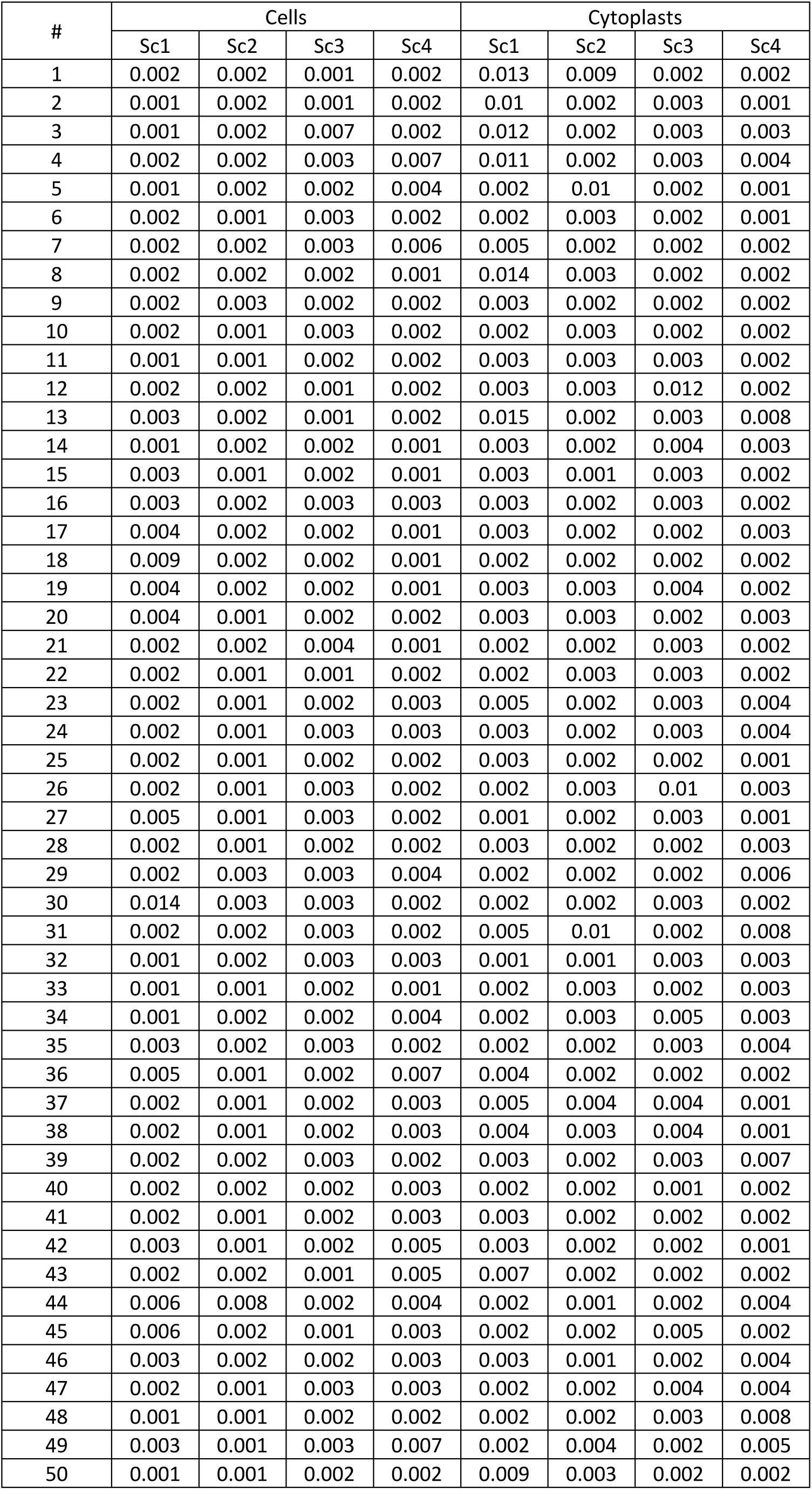
Approximate Entropy (ApEn) values for the 400 experimental amoeba trajectories.

| # | Cells |  |  |  | Cytoplasts |  |  |  |
| --- | --- | --- | --- | --- | --- | --- | --- | --- |
|  | Sc1 | Sc2 | Sc3 | Sc4 | Sc1 | Sc2 | Sc3 | Sc4 |
| 1 | 0.002 | 0.002 | 0.001 | 0.002 | 0.013 | 0.009 | 0.002 | 0.002 |
| 2 | 0.001 | 0.002 | 0.001 | 0.002 | 0.01 | 0.002 | 0.003 | 0.001 |
| 3 | 0.001 | 0.002 | 0.007 | 0.002 | 0.012 | 0.002 | 0.003 | 0.003 |
| 4 | 0.002 | 0.002 | 0.003 | 0.007 | 0.011 | 0.002 | 0.003 | 0.004 |
| 5 | 0.001 | 0.002 | 0.002 | 0.004 | 0.002 | 0.01 | 0.002 | 0.001 |
| 6 | 0.002 | 0.001 | 0.003 | 0.002 | 0.002 | 0.003 | 0.002 | 0.001 |
| 7 | 0.002 | 0.002 | 0.003 | 0.006 | 0.005 | 0.002 | 0.002 | 0.002 |
| 8 | 0.002 | 0.002 | 0.002 | 0.001 | 0.014 | 0.003 | 0.002 | 0.002 |
| 9 | 0.002 | 0.003 | 0.002 | 0.002 | 0.003 | 0.002 | 0.002 | 0.002 |
| 10 | 0.002 | 0.001 | 0.003 | 0.002 | 0.002 | 0.003 | 0.002 | 0.002 |
| 11 | 0.001 | 0.001 | 0.002 | 0.002 | 0.003 | 0.003 | 0.003 | 0.002 |
| 12 | 0.002 | 0.002 | 0.001 | 0.002 | 0.003 | 0.003 | 0.012 | 0.002 |
| 13 | 0.003 | 0.002 | 0.001 | 0.002 | 0.015 | 0.002 | 0.003 | 0.008 |
| 14 | 0.001 | 0.002 | 0.002 | 0.001 | 0.003 | 0.002 | 0.004 | 0.003 |
| 15 | 0.003 | 0.001 | 0.002 | 0.001 | 0.003 | 0.001 | 0.003 | 0.002 |
| 16 | 0.003 | 0.002 | 0.003 | 0.003 | 0.003 | 0.002 | 0.003 | 0.002 |
| 17 | 0.004 | 0.002 | 0.002 | 0.001 | 0.003 | 0.002 | 0.002 | 0.003 |
| 18 | 0.009 | 0.002 | 0.002 | 0.001 | 0.002 | 0.002 | 0.002 | 0.002 |
| 19 | 0.004 | 0.002 | 0.002 | 0.001 | 0.003 | 0.003 | 0.004 | 0.002 |
| 20 | 0.004 | 0.001 | 0.002 | 0.002 | 0.003 | 0.003 | 0.002 | 0.003 |
| 21 | 0.002 | 0.002 | 0.004 | 0.001 | 0.002 | 0.002 | 0.003 | 0.002 |
| 22 | 0.002 | 0.001 | 0.001 | 0.002 | 0.002 | 0.003 | 0.003 | 0.002 |
| 23 | 0.002 | 0.001 | 0.002 | 0.003 | 0.005 | 0.002 | 0.003 | 0.004 |
| 24 | 0.002 | 0.001 | 0.003 | 0.003 | 0.003 | 0.002 | 0.003 | 0.004 |
| 25 | 0.002 | 0.001 | 0.002 | 0.002 | 0.003 | 0.002 | 0.002 | 0.001 |
| 26 | 0.002 | 0.001 | 0.003 | 0.002 | 0.002 | 0.003 | 0.01 | 0.003 |
| 27 | 0.005 | 0.001 | 0.003 | 0.002 | 0.001 | 0.002 | 0.003 | 0.001 |
| 28 | 0.002 | 0.001 | 0.002 | 0.002 | 0.003 | 0.002 | 0.002 | 0.003 |
| 29 | 0.002 | 0.003 | 0.003 | 0.004 | 0.002 | 0.002 | 0.002 | 0.006 |
| 30 | 0.014 | 0.003 | 0.003 | 0.002 | 0.002 | 0.002 | 0.003 | 0.002 |
| 31 | 0.002 | 0.002 | 0.003 | 0.002 | 0.005 | 0.01 | 0.002 | 0.008 |
| 32 | 0.001 | 0.002 | 0.003 | 0.003 | 0.001 | 0.001 | 0.003 | 0.003 |
| 33 | 0.001 | 0.001 | 0.002 | 0.001 | 0.002 | 0.003 | 0.002 | 0.003 |
| 34 | 0.001 | 0.002 | 0.002 | 0.004 | 0.002 | 0.003 | 0.005 | 0.003 |
| 35 | 0.003 | 0.002 | 0.003 | 0.002 | 0.002 | 0.002 | 0.003 | 0.004 |
| 36 | 0.005 | 0.001 | 0.002 | 0.007 | 0.004 | 0.002 | 0.002 | 0.002 |
| 37 | 0.002 | 0.001 | 0.002 | 0.003 | 0.005 | 0.004 | 0.004 | 0.001 |
| 38 | 0.002 | 0.001 | 0.002 | 0.003 | 0.004 | 0.003 | 0.004 | 0.001 |
| 39 | 0.002 | 0.002 | 0.003 | 0.002 | 0.003 | 0.002 | 0.003 | 0.007 |
| 40 | 0.002 | 0.002 | 0.002 | 0.003 | 0.002 | 0.002 | 0.001 | 0.002 |
| 41 | 0.002 | 0.001 | 0.002 | 0.003 | 0.003 | 0.002 | 0.002 | 0.002 |
| 42 | 0.003 | 0.001 | 0.002 | 0.005 | 0.003 | 0.002 | 0.002 | 0.001 |
| 43 | 0.002 | 0.002 | 0.001 | 0.005 | 0.007 | 0.002 | 0.002 | 0.002 |
| 44 | 0.006 | 0.008 | 0.002 | 0.004 | 0.002 | 0.001 | 0.002 | 0.004 |
| 45 | 0.006 | 0.002 | 0.001 | 0.003 | 0.002 | 0.002 | 0.005 | 0.002 |
| 46 | 0.003 | 0.002 | 0.002 | 0.003 | 0.003 | 0.001 | 0.002 | 0.004 |
| 47 | 0.002 | 0.001 | 0.003 | 0.003 | 0.002 | 0.002 | 0.004 | 0.004 |
| 48 | 0.001 | 0.001 | 0.002 | 0.002 | 0.002 | 0.002 | 0.003 | 0.008 |
| 49 | 0.003 | 0.001 | 0.003 | 0.007 | 0.002 | 0.004 | 0.002 | 0.005 |
| 50 | 0.001 | 0.001 | 0.002 | 0.002 | 0.009 | 0.003 | 0.002 | 0.002 |

**Table S7.** Approximate Entropy (ApEn) values for the 400 shufled amoeba trajectories.

| # | Cells |  |  |  | Cytoplasts |  |  |  |
| --- | --- | --- | --- | --- | --- | --- | --- | --- |
|  | Sc1 | Sc2 | Sc3 | Sc4 | Sc1 | Sc2 | Sc3 | Sc4 |
| 1 | 2.09 | 2.067 | 2.088 | 1.9 | 1.942 | 2.109 | 2.062 | 1.893 |
| 2 | 2.081 | 2.091 | 2.047 | 1.924 | 2.013 | 1.858 | 1.735 | 2.058 |
| 3 | 2.005 | 2.081 | 1.945 | 2.018 | 2.122 | 1.909 | 1.965 | 1.94 |
| 4 | 2.029 | 2.074 | 1.955 | 1.943 | 1.759 | 1.964 | 1.946 | 1.941 |
| 5 | 1.941 | 1.926 | 1.716 | 1.957 | 1.878 | 1.96 | 1.664 | 2.07 |
| 6 | 1.976 | 2.05 | 1.881 | 1.916 | 1.478 | 1.957 | 2.069 | 2.053 |
| 7 | 2.081 | 2.042 | 1.961 | 2.053 | 1.702 | 1.849 | 1.701 | 1.993 |
| 8 | 2.105 | 1.932 | 2.044 | 1.923 | 1.98 | 1.793 | 1.681 | 1.79 |
| 9 | 2.072 | 1.725 | 2.03 | 1.792 | 1.905 | 2.003 | 1.577 | 2.017 |
| 10 | 1.878 | 2.092 | 1.922 | 2.051 | 2.07 | 1.945 | 1.826 | 2.02 |
| 11 | 2.079 | 2.087 | 2.109 | 1.919 | 1.914 | 1.752 | 1.736 | 1.995 |
| 12 | 2.084 | 2.083 | 2.104 | 2.02 | 1.909 | 1.725 | 2.037 | 1.786 |
| 13 | 1.92 | 2.108 | 2.075 | 2.004 | 1.921 | 1.532 | 1.683 | 1.743 |
| 14 | 2.062 | 2.021 | 1.916 | 2.028 | 1.298 | 1.83 | 1.704 | 1.975 |
| 15 | 2.103 | 2.094 | 1.553 | 1.967 | 1.825 | 1.937 | 1.824 | 1.813 |
| 16 | 2.027 | 2.033 | 1.997 | 1.75 | 1.811 | 2.072 | 1.966 | 1.793 |
| 17 | 1.92 | 2.128 | 1.945 | 2.08 | 1.943 | 1.893 | 2.035 | 1.825 |
| 18 | 1.99 | 1.853 | 1.872 | 1.967 | 1.824 | 1.691 | 1.846 | 1.856 |
| 19 | 1.612 | 2.09 | 2.063 | 2.104 | 1.985 | 1.69 | 1.635 | 2.017 |
| 20 | 2.022 | 2.066 | 2.081 | 1.854 | 1.618 | 1.818 | 1.802 | 1.724 |
| 21 | 1.943 | 2.055 | 1.912 | 2.086 | 1.959 | 1.954 | 1.948 | 1.951 |
| 22 | 1.679 | 2.092 | 2.067 | 2.044 | 1.969 | 1.837 | 1.744 | 1.601 |
| 23 | 1.966 | 2.065 | 2.089 | 1.967 | 1.725 | 1.772 | 1.851 | 1.896 |
| 24 | 1.742 | 2.042 | 1.997 | 1.825 | 1.69 | 2.019 | 1.825 | 1.471 |
| 25 | 1.925 | 2.023 | 2.067 | 1.511 | 1.731 | 2.04 | 1.682 | 1.996 |
| 26 | 1.946 | 2.044 | 1.998 | 1.921 | 1.844 | 1.804 | 1.881 | 1.763 |
| 27 | 2.029 | 1.945 | 1.937 | 1.986 | 2.1 | 1.948 | 1.682 | 2.018 |
| 28 | 2.099 | 2.055 | 1.996 | 1.804 | 2.059 | 2.015 | 1.807 | 1.929 |
| 29 | 2.017 | 2.047 | 2.055 | 1.797 | 1.809 | 1.843 | 1.914 | 2.029 |
| 30 | 2.016 | 2.066 | 2.104 | 1.95 | 2.021 | 1.872 | 1.812 | 1.648 |
| 31 | 1.95 | 2.083 | 2.034 | 1.778 | 1.619 | 1.78 | 1.932 | 1.944 |
| 32 | 2.077 | 1.998 | 1.934 | 2.014 | 1.993 | 2.074 | 2.01 | 1.933 |
| 33 | 1.99 | 1.754 | 1.98 | 2.1 | 1.979 | 1.914 | 1.959 | 2.022 |
| 34 | 1.9 | 2.091 | 1.907 | 1.929 | 2.078 | 1.906 | 1.9 | 1.688 |
| 35 | 1.927 | 1.853 | 1.926 | 2.017 | 1.934 | 1.631 | 1.991 | 1.962 |
| 36 | 1.993 | 2.111 | 1.895 | 1.964 | 1.927 | 2.061 | 2.1 | 2.035 |
| 37 | 2.013 | 2.081 | 1.964 | 1.483 | 1.957 | 1.583 | 1.803 | 2.074 |
| 38 | 2.039 | 2.013 | 2.066 | 1.967 | 1.451 | 1.825 | 1.575 | 1.996 |
| 39 | 1.907 | 1.992 | 1.808 | 1.892 | 1.927 | 1.769 | 1.915 | 1.843 |
| 40 | 1.961 | 2.04 | 1.944 | 1.908 | 1.966 | 1.95 | 2.089 | 1.71 |
| 41 | 2.056 | 2.084 | 2.023 | 1.814 | 1.99 | 1.969 | 1.943 | 2.092 |
| 42 | 1.913 | 2.069 | 2.077 | 1.761 | 1.819 | 1.764 | 2.065 | 2.002 |
| 43 | 2.046 | 2.058 | 2.098 | 1.97 | 1.963 | 2.005 | 2.045 | 2.037 |
| 44 | 1.699 | 1.901 | 1.899 | 1.871 | 2.072 | 2.047 | 2.013 | 1.832 |
| 45 | 1.855 | 1.999 | 1.982 | 2.039 | 1.931 | 1.992 | 1.712 | 1.774 |
| 46 | 2.059 | 2.08 | 2.104 | 1.991 | 1.618 | 1.964 | 2.06 | 1.801 |
| 47 | 1.627 | 2.078 | 1.887 | 1.606 | 1.923 | 2.073 | 1.915 | 1.791 |
| 48 | 1.794 | 2.114 | 2.029 | 1.942 | 2.062 | 2.027 | 1.842 | 1.947 |
| 49 | 2.011 | 2.095 | 1.877 | 1.99 | 2.032 | 2.076 | 1.979 | 1.449 |
| 50 | 2.07 | 2.07 | 1.895 | 1.835 | 1.95 | 1.89 | 1.888 | 1.582 |

**Table S8.**
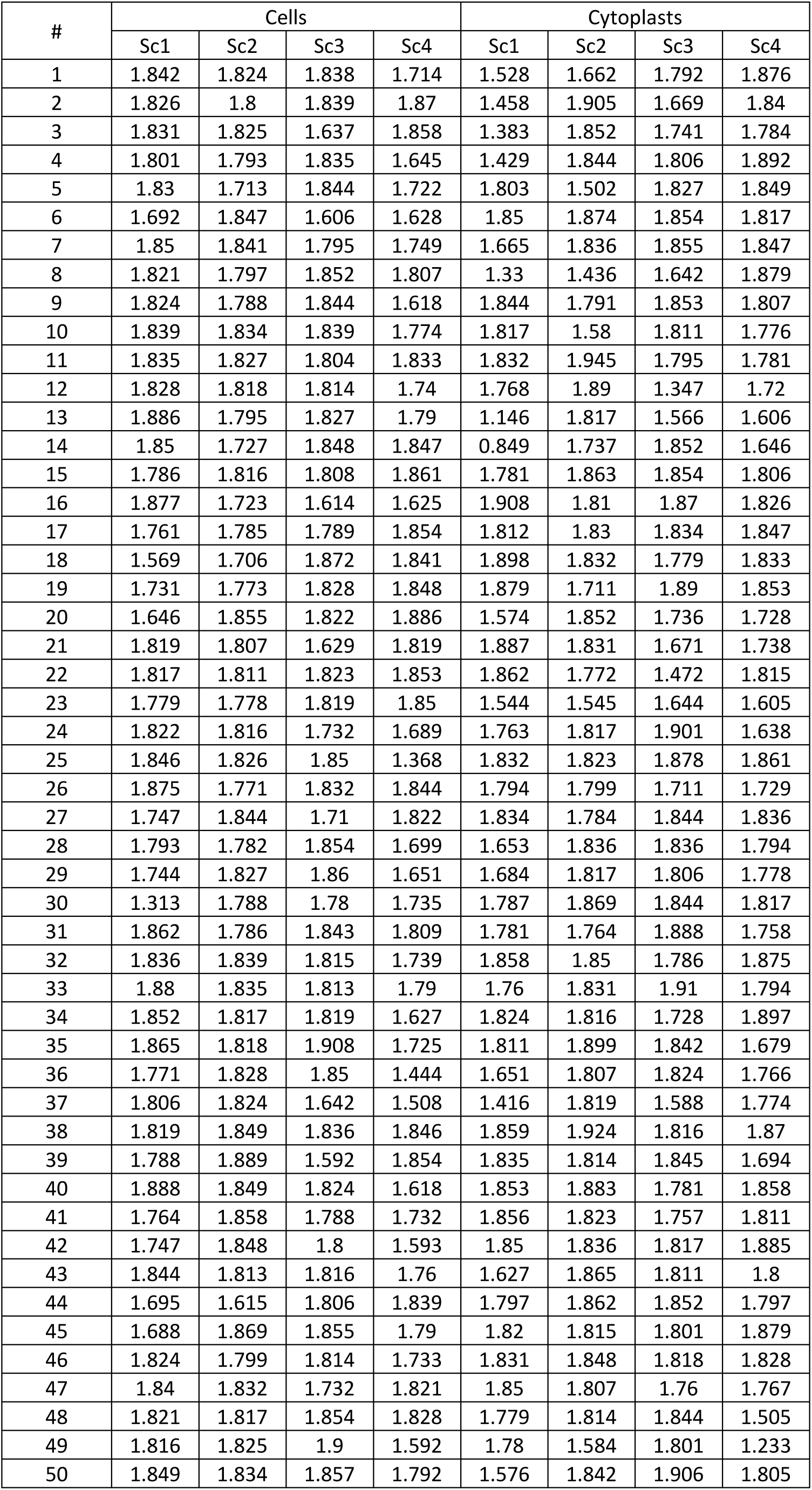
Detrended Fluctuation Analysis (DFA) scaling exponent γ values for the 400 experimental amoeba trajectories.

**Table S9.** Detrended Fluctuation Analysis (DFA) scaling exponent γ values for the 400 shufled amoeba trajectories.

| # | Cells |  |  |  | Cytoplasts |  |  |  |
| --- | --- | --- | --- | --- | --- | --- | --- | --- |
|  | Sc1 | Sc2 | Sc3 | Sc4 | Sc1 | Sc2 | Sc3 | Sc4 |
| 1 | 0.425 | 0.482 | 0.507 | 0.481 | 0.347 | 0.539 | 0.529 | 0.302 |
| 2 | 0.352 | 0.509 | 0.516 | 0.513 | 0.505 | 0.35 | 0.421 | 0.576 |
| 3 | 0.415 | 0.401 | 0.312 | 0.555 | 0.42 | 0.569 | 0.303 | 0.55 |
| 4 | 0.6 | 0.527 | 0.414 | 0.429 | 0.303 | 0.546 | 0.461 | 0.524 |
| 5 | 0.551 | 0.304 | 0.503 | 0.421 | 0.375 | 0.619 | 0.557 | 0.456 |
| 6 | 0.526 | 0.355 | 0.384 | 0.518 | 0.484 | 0.303 | 0.51 | 0.576 |
| 7 | 0.445 | 0.566 | 0.421 | 0.46 | 0.475 | 0.544 | 0.377 | 0.509 |
| 8 | 0.524 | 0.549 | 0.442 | 0.455 | 0.529 | 0.604 | 0.554 | 0.282 |
| 9 | 0.355 | 0.663 | 0.608 | 0.479 | 0.443 | 0.41 | 0.551 | 0.281 |
| 10 | 0.508 | 0.511 | 0.476 | 0.461 | 0.592 | 0.45 | 0.551 | 0.507 |
| 11 | 0.332 | 0.305 | 0.408 | 0.482 | 0.505 | 0.608 | 0.427 | 0.513 |
| 12 | 0.539 | 0.495 | 0.422 | 0.466 | 0.558 | 0.472 | 0.603 | 0.539 |
| 13 | 0.466 | 0.496 | 0.33 | 0.354 | 0.552 | 0.562 | 0.589 | 0.584 |
| 14 | 0.53 | 0.445 | 0.422 | 0.596 | 0.657 | 0.505 | 0.496 | 0.419 |
| 15 | 0.467 | 0.434 | 0.497 | 0.361 | 0.611 | 0.454 | 0.547 | 0.366 |
| 16 | 0.456 | 0.467 | 0.389 | 0.344 | 0.421 | 0.472 | 0.664 | 0.406 |
| 17 | 0.276 | 0.569 | 0.397 | 0.39 | 0.469 | 0.418 | 0.496 | 0.53 |
| 18 | 0.62 | 0.634 | 0.554 | 0.478 | 0.602 | 0.516 | 0.325 | 0.473 |
| 19 | 0.537 | 0.471 | 0.38 | 0.507 | 0.419 | 0.48 | 0.484 | 0.487 |
| 20 | 0.462 | 0.416 | 0.531 | 0.364 | 0.533 | 0.646 | 0.35 | 0.537 |
| 21 | 0.629 | 0.429 | 0.387 | 0.467 | 0.694 | 0.573 | 0.455 | 0.46 |
| 22 | 0.315 | 0.597 | 0.428 | 0.353 | 0.343 | 0.444 | 0.548 | 0.36 |
| 23 | 0.317 | 0.476 | 0.445 | 0.344 | 0.388 | 0.52 | 0.534 | 0.689 |
| 24 | 0.553 | 0.563 | 0.445 | 0.41 | 0.346 | 0.529 | 0.572 | 0.632 |
| 25 | 0.5 | 0.492 | 0.585 | 0.488 | 0.632 | 0.485 | 0.579 | 0.271 |
| 26 | 0.433 | 0.403 | 0.484 | 0.54 | 0.39 | 0.34 | 0.533 | 0.539 |
| 27 | 0.504 | 0.548 | 0.447 | 0.395 | 0.443 | 0.458 | 0.419 | 0.432 |
| 28 | 0.25 | 0.395 | 0.342 | 0.565 | 0.574 | 0.659 | 0.383 | 0.629 |
| 29 | 0.522 | 0.462 | 0.349 | 0.332 | 0.43 | 0.431 | 0.565 | 0.401 |
| 30 | 0.603 | 0.567 | 0.653 | 0.65 | 0.511 | 0.396 | 0.46 | 0.264 |
| 31 | 0.366 | 0.575 | 0.615 | 0.623 | 0.346 | 0.464 | 0.455 | 0.437 |
| 32 | 0.468 | 0.521 | 0.496 | 0.512 | 0.33 | 0.453 | 0.436 | 0.438 |
| 33 | 0.466 | 0.53 | 0.426 | 0.326 | 0.53 | 0.425 | 0.396 | 0.671 |
| 34 | 0.343 | 0.44 | 0.623 | 0.395 | 0.531 | 0.456 | 0.359 | 0.533 |
| 35 | 0.436 | 0.568 | 0.421 | 0.727 | 0.473 | 0.59 | 0.593 | 0.453 |
| 36 | 0.487 | 0.554 | 0.626 | 0.506 | 0.302 | 0.719 | 0.413 | 0.452 |
| 37 | 0.437 | 0.498 | 0.515 | 0.505 | 0.486 | 0.549 | 0.579 | 0.414 |
| 38 | 0.423 | 0.623 | 0.388 | 0.474 | 0.603 | 0.607 | 0.632 | 0.51 |
| 39 | 0.65 | 0.472 | 0.544 | 0.545 | 0.494 | 0.433 | 0.401 | 0.343 |
| 40 | 0.521 | 0.352 | 0.573 | 0.445 | 0.395 | 0.408 | 0.416 | 0.355 |
| 41 | 0.485 | 0.522 | 0.416 | 0.621 | 0.509 | 0.502 | 0.43 | 0.451 |
| 42 | 0.448 | 0.394 | 0.509 | 0.339 | 0.483 | 0.43 | 0.512 | 0.469 |
| 43 | 0.462 | 0.511 | 0.471 | 0.572 | 0.507 | 0.399 | 0.322 | 0.431 |
| 44 | 0.593 | 0.436 | 0.386 | 0.563 | 0.439 | 0.404 | 0.435 | 0.431 |
| 45 | 0.332 | 0.66 | 0.421 | 0.58 | 0.509 | 0.535 | 0.409 | 0.494 |
| 46 | 0.471 | 0.538 | 0.473 | 0.469 | 0.507 | 0.381 | 0.479 | 0.502 |
| 47 | 0.477 | 0.343 | 0.563 | 0.529 | 0.617 | 0.479 | 0.33 | 0.495 |
| 48 | 0.576 | 0.449 | 0.425 | 0.415 | 0.415 | 0.488 | 0.658 | 0.484 |
| 49 | 0.495 | 0.355 | 0.524 | 0.558 | 0.495 | 0.46 | 0.5 | 0.628 |
| 50 | 0.547 | 0.539 | 0.457 | 0.388 | 0.573 | 0.487 | 0.506 | 0.465 |

**Table S10.** Kruskal-Wallis and Wilcoxon comparison p-values to evaluate inter-cell-type and inter-scenario variability in kinetic and systemic properties.

| Cell type | Scenario | Intensity of Response | Directionality Ratio | Average Speed | RMSFα | RMSF Correlation time | MSDβ | ApEn | DFAy |
| --- | --- | --- | --- | --- | --- | --- | --- | --- | --- |
| Cells | Sc1-Sc2 | $10^{-4}$ | $10^{-5}$ | 0.022 | 0.007 | 0.992 | 0.078 | 0.003 | 0.717 |
|  | Sc1-Sc3 | 0.556 | 0.627 | 0.791 | 0.058 | 0.261 | 0.791 | 0.319 | 0.828 |
| | Sc1-Sc4 | 0.002 | 0.082 | $10^{-5}$ | 0.528 | 0.857 | 0.013 | 0.281 | 0.026 |
| | Sc2-Sc3 | 0.006 | $10^{-6}$ | 0.147 | 0.229 | 0.22 | 0.02 | $10^{-6}$ | 0.426 |
| | Sc2-Sc4 | $10^{-9}$ | $10^{-9}$ | $10^{-9}$ | 0.012 | 0.978 | $10^{-6}$ | $10^{-5}$ | 0.028 |
| | Sc3-Sc4 | $10^{-4}$ | 0.149 | $10^{-5}$ | 0.127 | 0.16 | 0.008 | 0.692 | 0.022 |
| | All | $10^{-9}$ | $10^{-9}$ | $10^{-8}$ | 0.013 | 0.493 | $10^{-4}$ | $10^{-5}$ | 0.048 |
| Cytoplasts | Sc1-Sc2 | 0.333 | 0.007 | 0.319 | 0.77 | 0.152 | 0.196 | $10^{-4}$ | 0.024 |
|  | Sc1-Sc3 | 0.652 | 0.009 | 0.662 | 0.807 | 0.931 | 0.038 | 0.068 | 0.165 |
|  | Sc1-Sc4 | 0.055 | 0.115 | 0.035 | 0.272 | 0.653 | 0.546 | 0.033 | 0.343 |
|  | Sc2-Sc3 | 0.497 | 0.828 | 0.169 | 0.92 | 0.099 | 0.333 | 0.051 | 0.354 |
|  | Sc2-Sc4 | 0.281 | 0.35 | 0.001 | 0.41 | 0.034 | 0.309 | 0.488 | 0.125 |
|  | Sc3-Sc4 | 0.073 | 0.224 | 0.065 | 0.497 | 0.678 | 0.071 | 0.467 | 0.637 |
|  | All | 0.182 | 0.023 | 0.013 | 0.738 | 0.183 | 0.122 | 0.007 | 0.126 |
| Cells Vs. Cytoplasts | Sc1 | $10^{-4}$ | 0.033 | $10^{-4}$ | 0.068 | 0.074 | 0.02 | $10^{-4}$ | 0.112 |
| | Sc2 | $10^{-9}$ | $10^{-5}$ | $10^{-9}$ | $10^{-7}$ | 0.445 | $10^{-4}$ | $10^{-6}$ | 0.251 |
| | Sc3 | $10^{-6}$ | 0.309 | 0.001 | $10^{-5}$ | 0.238 | 0.749 | 0.088 | 0.622 |
|  | Sc4 | 0.488 | 0.368 | 0.017 | 0.035 | 0.004 | 0.398 | 0.588 | 0.112 |
| | Sc1-Sc2 | 0.001 | 0.743 | $10^{-6}$ | 0.111 | 0.582 | 0.163 | 0.316 | 0.488 |
| | Sc1-Sc3 | $10^{-4}$ | 0.565 | 0.002 | 0.088 | 0.039 | 0.603 | 0.014 | 0.801 |
|  | Sc1-Sc4 | 0.007 | 0.642 | 0.12 | 0.333 | 0.019 | 0.046 | 0.143 | 0.414 |
| | Sc2-Sc3 | $10^{-11}$ | $10^{-4}$ | $10^{-6}$ | $10^{-6}$ | 0.035 | 0.013 | $10^{-10}$ | 0.942 |
| | Sc2-Sc4 | $10^{-8}$ | $10^{-6}$ | 0.001 | $10^{-5}$ | 0.019 | $10^{-5}$ | $10^{-5}$ | 0.395 |
| | Sc3-Sc4 | 0.002 | 0.712 | 0.123 | $10^{-4}$ | 0.122 | 0.04 | 0.391 | 0.281 |
| | All | $10^{-12}$ | 0.047 | $10^{-8}$ | $10^{-11}$ | 0.001 | 0.011 | $10^{-7}$ | 0.993 |

